# Multi-scale characterisation of homologous recombination deficiency in breast cancer

**DOI:** 10.1101/2023.08.23.554414

**Authors:** Daniel H. Jacobson, Shi Pan, Jasmin Fisher, Maria Secrier

## Abstract

**Background:** Homologous recombination is a robust, broadly error-free mechanism of double strand break repair, and deficiencies cause reliance on alternative repair processes, leading to sensitivity to PARP inhibitors. Patients displaying homologous recombination deficiency can be identified using ‘mutational signatures’. However, these patterns are difficult to reliably infer from exome sequencing. Additionally, as mutational signatures are a historical record of mutagenic processes, this limits their utility in describing the current status of a tumour.

**Results:** Here, we apply two methods for classifying homologous recombination deficiency in breast cancer to explore the features and heterogeneity associated with this phenotype. Firstly, we apply a likelihood-based method which leverages small insertions and deletions to improve classification of homologous recombination deficiency in exome sequenced breast cancers. We find that whilst BRCA+ homologous recombination deficient samples display strong similarities to those harbouring BRCA defects, they deviate in microenvironmental features such as hypoxic signalling. Secondly, using this classification we develop a 228-gene transcriptional signature which simultaneously characterises homologous recombination deficiency and BRCA1/2-defect status, and is linked with PARP inhibitor response. Finally, we apply this signature to single-cell RNA-sequenced breast cancer cohorts to study homologous recombination deficiency at single cell resolution and demonstrate that these cells present a distinct milieu of interactions with immune cells in their microenvironment compared to their HR proficient counterparts, which could inform checkpoint blockade for HRD tumours.

**Conclusions:** We apply multi-scale approaches to characterise homologous recombination deficiency in breast cancer through the development of mutational and transcriptional signatures. We show that indels, even at low levels, can improve homologous recombination deficiency classification. Additionally, we demonstrate the heterogeneity of homologous recombination deficiency, especially in relation to BRCA status, and show that indications of this feature can be captured at a single-cell level, enabling further investigations into interactions between DNA repair deficient cells and their tumour microenvironment.

## BACKGROUND

Maintaining genomic integrity is an essential process for ensuring the sustained survival of cancer cells and is enabled via a complex network of pathways forming the DNA damage response (DDR)(1,2). Homologous recombination (HR) is a robust method of repairing double strand breaks, and HR deficiency (HRD) causes the cell to become dependent on alternative repair processes such as non-homologous end joining (NHEJ) and theta-mediated end joining (TMEJ)(3). These reliances provide therapeutic opportunities to target HRD cells with treatments such as poly ADP-ribose polymerase (PARP) and polymerase θ (Polθ) inhibitors(4–7). PARP inhibitors have already been accepted for clinical use for patients with breast and ovarian cancer harbouring mutations in *BRCA1* and *BRCA2*(8–10). However, a significant proportion of cancer patients appear to display evidence of HRD without harbouring these biomarkers(11–13). Consequently, identifying patients demonstrating HRD, who may therefore benefit from these therapies, has received close attention.

One method for identifying HRD is using patterns of genomic aberrations known as ‘mutational signatures’(14). These signatures act as markers of prior mutagenic events and features, and identifying them within the cancer genome can highlight the driving forces behind the development of a given tumour. Signatures of HRD were uncovered as early as initial studies of mutational signatures(15), and HRD has since been linked with signatures of single-base substitutions (SBS), small insertions and deletions (indels), and structural variants(16,17). Specific classifiers have also been created involving the integration of multiple signatures(11) and individual mutational events(12). However, these classifiers work most optimally when applied to whole genome sequenced (WGS) data, and do not perform as well when capturing a reduced representation of the genome e.g. through exome sequencing, which is a common feature of large-scale cancer genomics datasets such as The Cancer Genome Atlas (TCGA).

Alternatives have included measures of chromosomal instability such as copy number signatures, which have been identified in a range of cancers(18–21) as well as the Myriad HRD index score(22,23), which integrates multiple large-scale genomic events to provide a broader measure of HRD(24–26). The Signature Multivariate Analysis (SigMA) computational tool applied a likelihood approach to classify both exomes and targeted panel sequenced data as SBS3-enriched(27), which has been shown to successfully predict olaparib response in breast and ovarian cancers(28). However, the scope of SigMA has been limited to the SBS3 signature alone as an HRD marker, and the inclusion of HRD-associated indel events offers potential for improved classification.

Whilst mutational signatures of HRD have shown substantial potential for predicting PARP inhibitor sensitivity, one drawback of this method is that, because these signatures result from the accumulation of mutations induced by various carcinogens acting from the very early stages of cancer initiation, they are by definition a feature of the history of a tumour, as opposed to its current state. This is of particular importance when it comes to HRD, as mechanisms of HR revival, such as BRCA1 reversion mutations and loss of 53BP1, lead to the emergence of PARP inhibitor resistance in BRCA1-defective patients(29,30). A signature based on the expression profile of a sample would, therefore, be more adept at describing the most recent state of a tumour. Transcriptional signatures have been applied to characterise various features associated with HRD, including BRCA1 loss in patients(31), HR gene knockdown in cell lines(32), PARP inhibitor sensitivity(33), and chromosomal instability(34). Additionally, the presence of signature SBS3 has been used to develop a gene signature for HRD classification in TNBC(35). This disregards the divergent consequences of BRCA1 and BRCA2 defects due to their different roles in governing HR function: whilst BRCA2 is intrinsically involved in HR via the recruitment of RAD51 to exposed single-stranded DNA, BRCA1 functions upstream of HR and is involved in determining the choice of repair pathway by inhibiting 53BP1, which drives end protection and therefore NHEJ(36,37). A transcriptional signature reflecting this heterogeneity may prevent a skewness towards BRCA1-type HRD, whilst also shedding light on the emergence of HRD in BRCA-positive patients.

In order to perform multi-scale characterisation and explore the heterogeneity of HRD, we present a mutational signature-based classifier of HRD for exome sequenced breast cancers which we then apply to develop a transcriptional signature of HRD and BRCA1/2 deficiency. We demonstrate that leveraging HRD-associated indel events improves HRD classification in downsampled WGS samples and in characterising BRCA-defective samples from the TCGA-BRCA cohort as HRD. Additionally, whilst BRCA+ and BRCA-defective HRD samples are broadly similar regarding standard hallmarks of HRD, BRCA+ samples show deviations in mutational and phenotypic features such as a comparative decrease in hypoxia. Using matched RNA-sequencing data, we then employ multinomial elastic net logistic regression to develop a 228-gene transcriptional signature which can be used to simultaneously predict BRCA1/2-deficiency and HRD status, and is linked with response to PARP inhibitors in both cell lines and patients from the I-SPY2 trial(38). Finally, we apply the signature to explore HRD at the cellular level from single cell RNA-sequencing data, and demonstrate substantial deviations in patterns of crosstalk with the tumour microenvironment (TME) between HRD and HR-proficient tumour cells. Together, these findings demonstrate the value of multi-scale examination of complex phenotypes like HRD and offer opportunities to improve research into the causes and consequences of such deficiencies in human cancers.

## RESULTS

### Establishing HRD-associated signature phenotypes in whole genome sequenced breast cancers

Prior to studying HRD in exome sequenced samples, we aimed to develop an initial understanding of the recurring profiles of mutational signatures seen in breast cancers. This was achieved by calculating the contributions of breast cancer-associated SBS and indel signatures in 614 whole genome sequenced breast cancers obtained from the International Cancer Genome Consortium (ICGC), and then clustering these profiles to established ‘signature phenotypes’. According to finite mixture modelling, the ICGC cohort could be grouped into 20 clusters, seven of which were assigned as ‘HRD’ due to the enrichment of SBS3 and the indel signatures ID6 and ID8 (Supplementary Figure 1a-b; Supplementary Table 1).

As expected, *BRCA-*defective samples were strongly enriched within the seven HRD clusters, as were samples labelled as HRD by either HRDetect(11) or CHORD(12), with all BRCA-defective samples appearing in an HRD cluster (Supplementary Figure 1c). 117/120 (97.5%) of samples with HRDetect scores greater than 0.7 appear in an HRD-associated cluster. Considering samples with known BRCA status, 75/135 (55.6%) of samples within the HRD clusters are BRCA-defective, similar to the 74/120 (61.7%) of samples with HRDetect scores greater than 0.7. This is demonstrative of the fact that, even accounting for defects in a range of HR genes such as *RAD51C* and *PALB2* which would increase the precision of these classifiers, the source of HRD for many samples is unknown, and therefore the specificity for any HRD classifier is not expected to reach 100%.

Interestingly, BRCA1 and BRCA2-defective samples could also be broadly separated. Whilst this has been demonstrated using CHORD(12), we show that it can also be achieved using mutational signatures. BRCA2-defective samples are enriched in clusters characterised by increased contribution of the ID6 signature, referred here as ‘BRCA2-type HRD’. The ID6 signature displays deletions at microhomologous regions flanking double strand breaks, indicative of high TMEJ activity(39). Alternatively, BRCA1-defective samples appear in clusters featuring increased ID8 signature contributions, which we call ‘BRCA1-type HRD’, associated with non-homologous end joining(17). Since BRCA1 is heavily involved in determining how a double strand break will be processed, BRCA1-defective samples will naturally rely on non-homologous end joining to be their primary method of DSB repair. Each type of HRD cluster shows high sensitivity for classifying their respective BRCA defect, with BRCA1 and BRCA2 defects being correctly classified with sensitivities of 68.9% and 70.0% respectively (Supplementary Figure 1d). Whilst CHORD shows greater specificity (73.3% and 93.3% respectively) there is concordance between the BRCA-type HRD clustering and CHORD classification (78.3% for BRCA1-type HRD and 72.2% for BRCA2-type HRD). Thus the indel signatures may not only be useful as HRD-associated signatures, but also shed light on the method employed by a sample for tolerating that type of HRD.

### Evaluating HRD in exome sequenced breast cancers

A minimum of 50 mutations are generally believed to be required for reliable mutational signature inference(40). By this criterion, SBS signature analysis is unsuitable in 558/968 whole-exome sequenced samples from the TCGA-BRCA cohort, and indel signature analysis is unsuitable in 958/968 (Supplementary Figure 2). In particular, these samples display a median indel load of three, and a mean of 5.39, with 94 samples harbouring zero indel events. There is an opportunity to overcome such limitations when it comes to identifying HRD in exome sequenced cancers by employing the previously described signature phenotypes in whole cancer genomes: rather than identifying the relevant signatures themselves, the signature profile of a cluster can be predicted instead. To achieve this, we developed a likelihood-based computational method which would enable the assignment to the 20 signature phenotypes without the need to calculate the prevalence of specific signatures (Figure 1a). For each of the 20 clusters, a mean mutational spectrum was calculated, representing a ‘ground-truth’/baseline profile against which new sample spectra can be compared (Supplementary Figure 3). Subsequently, each mutation in the profile of a new sample will influence the likelihood of that sample being assigned to each of the signature phenotypes, with the prior probabilities of assignment to each cluster determined by their size in the ICGC-BRCA cohort.

**Figure 1.**
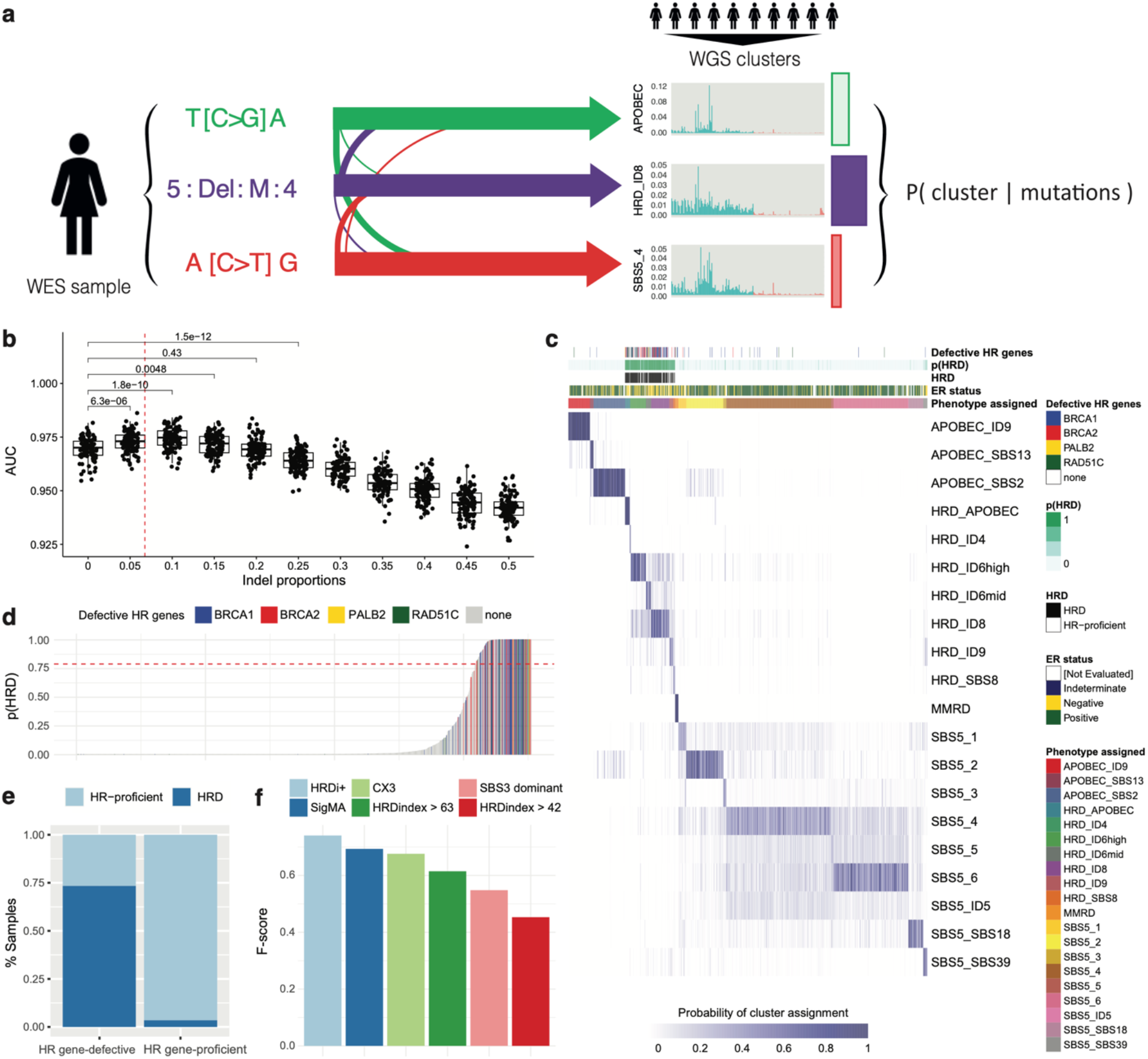
Evaluating HRD in exome sequenced breast cancers. (a) Workflow for HRD classification of an exome-sequenced breast cancer sample. Each sample contains a profile of mutations, each of which has a probabilistic association with each of the 20 signature phenotypes, defined by a representative signature profile inferred from WGS data. The mutational profile is collated to calculate the probability of assignment of the respective sample to each of the 20 clusters. (b) Simulation analysis of SBS3-enrichment classification of ICGC samples downsampled to 50 mutational events constrained to varying indel proportions. Adding a small percentage of indels is sufficient to improve classification. AUC = Area Under the ROC Curve for SBS3 enrichment classification. The dotted red line represents the mean proportion of indel events in the TCGA-BRCA cohort. (c) Classification of 968 exome sequenced breast cancer samples from TCGA. The heat map indicates the probability of each sample (column) being assigned to each signature phenotype (rows). Samples are annotated by ER status and HR gene defects. The value p(HRD) is the sum of probabilities of assignment across the seven HRD-associated phenotypes. The label ‘Phenotype assigned’ refers to the phenotype to which the respective sample has the highest probability of assignment. (d) Summary of HRD cluster assignment probabilities across the TCGA-BRCA cohort. Samples with a total probability of HRD assignment greater than 0.79 (as shown by the dotted red line) would be assigned as HRD, whereas the rest would be deemed HR-proficient. (e) HRD classification of HR gene-defective and -positive samples in the TCGA-BRCA cohort. ‘HRD’ refers to samples with a probability of HRD assignment greater than 0.79. (f) F-score comparisons of HRD classifiers for exomes. ‘HRDi+’ refers to the classifier developed in this study. The HRD index is presented using cut-offs of 42 and 63.

To initially demonstrate the capacity of this technique and the accuracy gained by considering indel events, we simulated exome sequenced samples by downsampling whole-genomes from ICGC data to see if our framework would correctly classify the downsampled data according to clusters defined by mutational signatures (see Methods). For these simulations, we based the clustering solely on SBS signatures as a conservative approach that would avoid any bias from indel events which might favour our methodology. We sampled events from each ICGC sample with replacement, constraining them to specific proportions of indel events to demonstrate how their inclusion improved classification. Simulated exomes were set to 25, 50, and 100 mutations. At mutation loads of 50, classification of SBS3 enrichment improved when 5-10% of the simulated mutations were constrained to indel events, indicating that even a small number of indels could improve HRD classification (Fig. 1b). As the simulated mutational load decreased from 100 to 25, a larger proportion of indels was required to enable substantial improvement over using SBS events alone (5% versus 20%, Supplementary Figure 4). The mean and median indel proportions across the TCGA-BRCA cohort are 6.75% and 5.88% respectively, thereby demonstrating that the results of these simulations align with real-world features and that the inclusion of indel events improves HRD classification.

Given the demonstrated and known importance of indel events for HRD classification, we sought to understand whether increasing the weights of indels within the likelihood distributions would improve classification. The above analysis was repeated in WGS samples downsampled to 50 mutations, and the respective indel proportions within the likelihood distributions were multiplied by factors increasing from 1/5 to 5 (see Methods). We demonstrate that whilst HRD classification is broadly unaffected when the indel likelihood weight is decreased, accuracy decreases significantly once it is increased, thereby showing that overestimating the importance of these indel events will hamper the accuracy of HRD classification (Supplementary Figure 5). Thus, from our analysis we conclude that the optimal HRD classification is attained when indel spectra are equally weighted as the SNV ones.

Finally, we asked if the full set of 20 phenotypes seen in WGS data could be captured at lower mutational loads. To this end, we conducted simulations to test the reclassification ability for each phenotype following subsampling (Supplementary Figure 6). This demonstrated that reclassification of specific APOBEC-enrichment clusters was highest at low mutational loads, followed by HRD-enrichment. Additionally, in cases where a sample was misclassified, it was most often assigned to a broader associated cluster. Thus the APOBEC and HRD phenotypes are reliably identified as such even when the mutation load in a tumour is low or not fully captured by the sequencing technique.

We next applied the likelihood-based classifier to 968 exome-sequenced breast cancer samples obtained from TCGA (Figure 1c; Supplementary Table 2). The HR gene-defective samples were mainly assigned across the seven HRD-associated clusters. The overall probability of HRD classification was calculated as the sum of probabilities of assignment to the seven HRD-associated clusters, and samples with a classification probability greater than 0.79 were labelled as HRD. This value was selected to maximise the accuracy of the classifier for identifying patients with defects or alterations in HR genes (BRCA1, BRCA2, RAD51C, and PALB2, hereafter termed “HR gene-defective”), as determined using an F-score (Methods; Supplementary Figure 7a). This method was selected to ensure sufficient sensitivity for classifying samples harbouring known HR gene defects, whilst maximising the confidence in HRD classification of HR gene-positive samples. Overall, HR gene-defective samples from TCGA were labelled as HRD with an AUC of 0.91 (Figure 1d, Supplementary Figure 7b), 73.4% sensitivity and 74.8% specificity (Figure 1e).

In general, the accuracy of the classifier is greater when considering ER-negative samples compared to ER-positive ones, likely due to the increased enrichment of HR gene-defective samples within ER-negative samples (31.6% versus 6.79%) (Supplementary Figure 7c-d, Supplementary Table 2). It is known that HRD features can vary substantially between cancer types(41), and therefore in some cases redefining specific thresholds based on a given context can improve classification. However, in both cases, the probability threshold of 0.79 performs well for maximising the resulting F-scores.

A similar performance was observed when applying the method in an independent exome-sequencing validation dataset of 186 breast cancer patients from the SMC Korean breast cancer cohort(42). Here, 13/20 (65%) of BRCA-defective patients were classified as HRD, increasing to 17/20 (85%) when applying a probability cut-off of 0.5 (Supplementary Figure 8).

Our method outperformed other signature-based methods such as SigMA(27), the CX3 copy number signature from Drews et al(20), or SBS3-based identification alone in terms of both specificity and sensitivity for classifying HR gene-defective samples as HRD (Figure 1f). It also outperformed the HRD index score, which is based on the levels of loss of heterozygosity, large scale transitions and telomeric allelic imbalances in a sample (Figure 1f). We note that the HRD large-scale genomic aberration features required for the Myriad HRD test are often imperfectly called from exome-sequencing data, and that efforts have been made to optimise the calling of these features(41). Therefore, we have used two different thresholds of HRD classification using this HRD index score based on literature-reported cutoffs(43,44). Whilst our classifier had a recall in TCGA of only 73.4%, generally lower than some alternative methods proposed, it demonstrated a far superior precision of 74.8%, demonstrating an increased stringency and confidence for HRD classification (Supplementary Figure 9).

Assignment to HRD clusters occurs more frequently for BRCA2-defective samples (93%) compared with BRCA1-defective (72%) and RAD51-defective (68%) samples (Supplementary Figure 10). However, whilst HRD classification was strong amongst HR gene-defective samples, unlike in the ICGC data, the classifier was unable to assign BRCA-defective samples to BRCA1- or BRCA2-type HRD clusters specifically, with respective sensitivities of 55.0% and 37.0% (Supplementary Figure 10). This is likely due to the scarcity of indel events appearing in exome-sequenced samples.

### Hallmarks of HRD

Primary TCGA-BRCA samples labelled as HRD according to the classifier displayed numerous hallmarks associated with DDR deficiencies (Figure 2), including greater levels of large-scale genomic scarring (Fig. 2a), and high levels of the CX3 copy number signature (Fig. 2b), which has been linked with impaired HRD alongside increased replication stress and impaired damage sensing(20), and which performed particularly well for HR gene-defect HRD classification (Supplementary Figure 11). HRD samples displayed increased expression of POLQ (Fig. 2c), indicating a greater reliance on theta-mediated end joining for double strand break repair(6,45,46), as well as increased proliferative capacity (Fig. 2d), which is expected to be associated with HRD(47).

**Figure 2.**
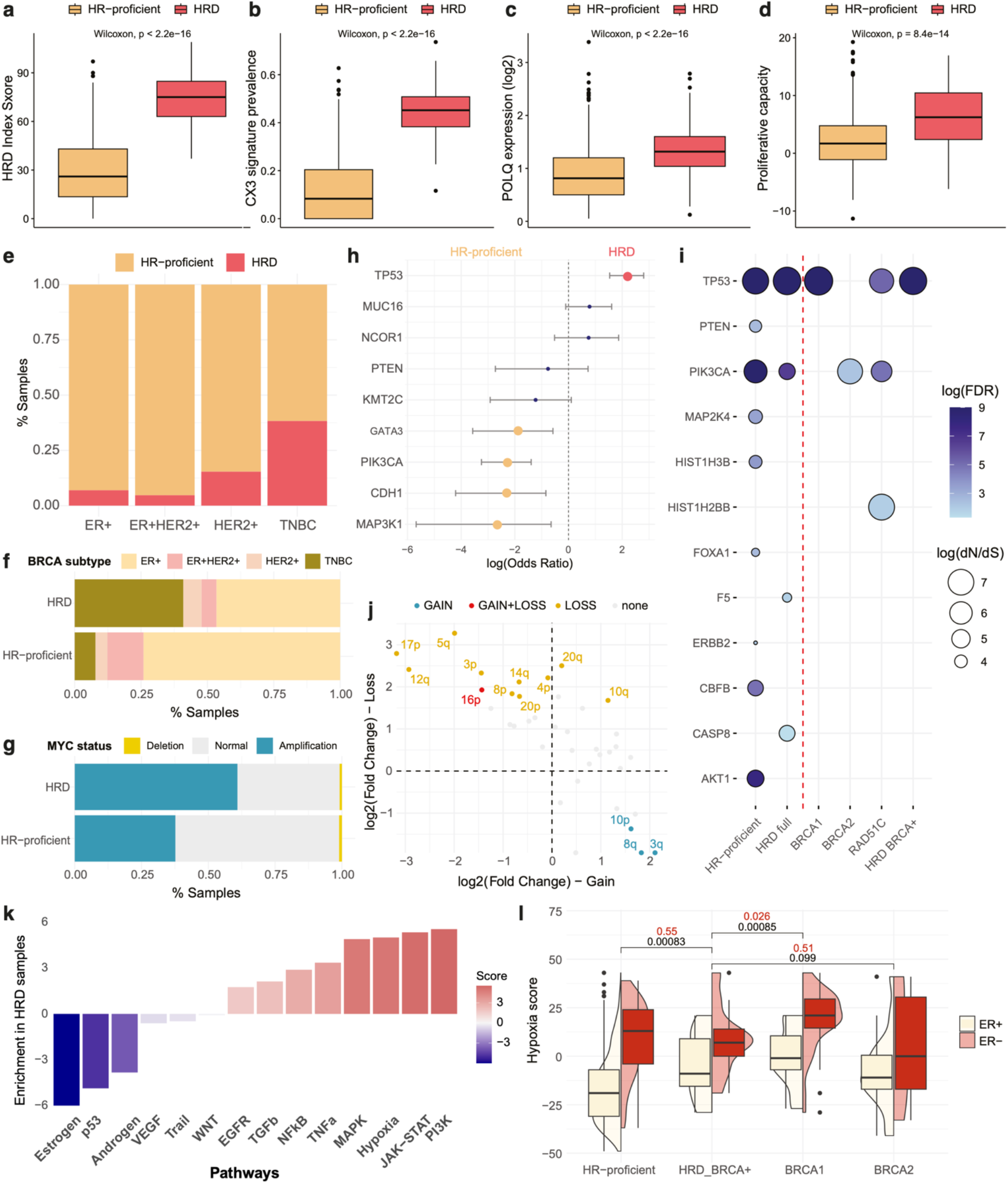
Genomic and transcriptional hallmarks of HRD. Association between HRD status and (a) the Myriad HRD index score, (b) contribution of the CX3 copy number signature, (c) POLQ expression, and (d) a transcriptional measurement of proliferation/cell cycle arrest capacity. (e) Enrichment of HRD across breast cancer subtypes. (f) Enrichment of breast cancer subtypes across HRD status. (g) Association between HRD status and amplification of MYC. (h) Enrichment/depletion of somatic nonsynonymous mutations in key cancer genes in HRD and HR-proficient breast cancer samples. A positive log(Odds Ratio) indicates enrichment in HRD samples. (i) Positive selection of cancer genes in HR-proficient, all HRD, BRCA1-defective, BRCA2-defective, RAD51C-defective, and HRD BRCA+ breast cancer samples. Circle size indicates the strength of positive selection according to the dN/dS ratio. (j) Comparison of chromosome arm loss and gain events between HRD and HR-proficient breast cancer samples. Positive values indicate enrichment in HRD against HR-proficient samples, whilst the x and y axes indicate enrichment for chromosome arm gains and losses respectively. (k) Results of differential pathway activity analysis between HRD and HR-proficient breast cancer samples across 14 signalling pathways ordered by the Normalised Enrichment Score (NES). Positive scores indicate pathway enrichment in HRD samples. (l) Comparison of hypoxia scores in the TCGA-BRCA cohort according to the Buffa transcriptional signature across HRD/BRCA-defect categories, split by ER status. P-values refer to Wilcoxon testing between each group and the HRD-BRCA+ group, tested across all samples (black) and ER-negative samples only (red).

Within TCGA, we see that HRD samples are significantly enriched amongst TNBC samples, with 38.3% of TNBC samples labelled as HRD (with TNBC constituting 40.9% of all HRD samples) compared with 7.1% amongst the remaining samples (Fig. 2e-f), which was also observed in the SMC breast cancer cohort(42). This aligns with prior information pointing to TNBC samples in TCGA being highly enriched for HR gene defects (43.2% compared with 7.2% amongst the remaining samples). We also classify 55% of the TNBC samples in the SMC cohort as HRD, which is aligned with their findings that 85% of these samples displayed at least some SBS3 signature contribution (Supplementary Figure 8c). Finally, we observed an association between HRD and amplification of MYC (Fig. 2g), which could imply an increase in replication stress(48).

HRD and HR-proficient breast cancers in TCGA were shown to display differential enrichment of mutational drivers of tumorigenesis (Fig. 2h). *TP53* mutations were significantly enriched in HRD samples, in agreement with their common co-occurrence with BRCA1/2 defects(49,50). In contrast, the genes *CDH1*, *MAP3K1*, *PIK3CA*, and *GATA3* were more frequently altered in HR-proficient samples. Mutations in *GATA3*, which is involved in normal mammary gland development and has been previously associated with ER-positivity, occur frequently in breast cancer, in particular frameshift indels(16,51,52), and were strongly enriched in a cohort of Nigerian breast cancer patients(53).

We applied the dN/dS method(54) to identify signals of positive selection for mutations within the HRD and HR-proficient groups (Fig. 2i). Unsurprisingly, mutations in *TP53* and *PIK3CA* were positively selected within both groups, acting as generic drivers in this cancer. Seven genes were positively selected in the HR-proficient group but not HRD. In contrast, *CASP8* and *F5* were positively selected only in HRD samples. Caspase signalling has previously been hypothesised as a driver of cell death following the induction of cGAS-STING and interferon signalling induced by knock-out of BRCA2, indicating that positive selection for *CASP8* mutation could allude to a method of maintaining cell viability in the context of increased chromosomal instablility(55).

BRCA1- and BRCA2-defective samples showed different patterns of positive selection according to dN/dS analysis, with BRCA1-defective and HRD BRCA+ samples showing positive selection only for *TP53*, whilst BRCA2-defective samples only demonstrated this in *PIK3CA* (Fig. 2i). (56) Due to the chromosomal instability associated with HRD, we also sought to highlight associated copy number aberrations. Chromosome arms were more likely to be enriched for losses than gains in HRD samples, aligning our knowledge of loss of heterozygosity as an HRD signature(26,56) (Fig 2j). Only chromosome arm 16p was significantly enriched for both losses in HRD samples, and gains in HR-proficient samples. Notably, the 16p arm carries the *PALB2* gene, which is strongly involved in HR and has been associated with PARPi sensitivity(57–59). Only three arms were significantly enriched for gains in HRD samples, which were 3q, which carries *POLQ*, 10p, which carries the *DCLRE1C* gene, which encodes for the Artemis protein that is essential for NHEJ activity(60), and 8q, which carries the *SPIDR* and *RAD54B* genes, encoding accessory factors for RAD51 activity during homologous recombination(61,62), indicating a potential compensatory mechanism in these patients.

Given the variation in mutation selection across HRD samples, we sought to further analyse how BRCA+ HRD samples compare to those harbouring BRCA defects. HRD-BRCA+ samples showed significantly increased levels of HRD hallmarks in comparison to HR-proficient samples (Supplementary Figure 11a). Whilst they showed a slight decrease in CX3 copy number signature contribution compared with BRCA-defective samples, these samples showed no difference in *POLQ* expression or proliferative capacity, indicating that they were displaying a clear HRD phenotype despite their BRCA+ status. Additionally, patients with an HRD classification probability between 0.5 and 0.79 display significantly lower levels of HRD hallmarks compared with those exceeding the threshold, further indicating increased confidence in our HRD classification (Supplementary Figure 11b).

Additionally, we checked the somatic mutation status of HRD HR gene-positive samples in comparison with those carrying HR gene defects. In commonly altered cancer genes, there were no discernible differences between BRCA+ and HR gene-defective samples (Supplementary Figure 12a). Of the 27 HRD HR gene-positive samples, 25 (93%) presented a non-silent mutation in a DDR gene, of which 10 (37%) carried mutations in a double strand repair gene, according to a curated list of genes associated with DDR(63). Whilst DDR gene mutations were similarly frequent in BRCA-defective HRD samples (96%), the proportion of double strand repair-mutated samples, excluding BRCA1 and BRCA2, was significantly higher at 61%. Within HRD BRCA+ samples, 17/27 (63%) carried a *TP53* mutation (similar to 68% amongst HR gene-defective samples), with the next most commonly mutated DDR genes being *ARID1A* and *MDM4*, a p53 inhibitor, although this was only in 2/27 cases each (Supplementary Figure 12b). Interestingly, mutations in *ARID1A* have previously been associated with PARPi sensitivity(64), indicating a potentially rare cause of HRD. The 16p chromosome arm, carrying the *PALB2* gene, was marginally enriched for gains amongst HRD HR gene-positive samples (Supplementary Figure 12c). However, following multiple testing correction, no genes were enriched for gains or losses within HRD samples depending on HR gene defects (Supplementary Figure 12d).

To gain a broader perspective on the differences driven by BRCA status in HRD samples, we analysed variation in pathway activity using decoupleR(65). Unsurprisingly, estrogen and p53 signaling were significantly downregulated in HRD samples (Fig. 2k). We also found that the hypoxia signalling response was substantially upregulated in HRD compared with HR-proficient samples (Fig. 2k). It has previously been demonstrated that BRCA-defective samples from TCGA display increased hypoxia scores compared with BRCA+ samples(66). Here, we found that HRD HR gene-positive samples also show increased hypoxia scores against HR-proficient samples, although this association disappears when analysing ER-negative samples alone (Fig. 2l). Interestingly, whilst these samples have lower hypoxia scores compared to BRCA1-defective samples, this was not observed in comparison to BRCA2-defective samples. A two-way ANOVA revealed that even after accounting for ER-status, both BRCA-defect (F(2,801) = 11.28, p = 1.5e-05) and HRD status (F(1,801) = 11.28, p = 8.2e-04) were significantly associated with hypoxia scores (Supplementary Table 3). Severe hypoxia has been shown to lead to PARPi sensitivity in HR-proficient tumours(67), and hypoxia has also been shown to inhibit ER expression in breast cancer cells(68), potentially explaining the similar hypoxia levels across BRCA+ ER-negative breast cancers. However, hypoxia as a BRCA-independent mechanism of HRD requires further analysis and experimental validation that is beyond the scope of this study.

Overall, we confirm that the samples classified as HRD via our signature-based method display numerous HRD-associated hallmarks, and demonstrate that HRD and HR-proficient samples show noticeable variation in genomic profiles. Additionally, HRD samples can show variation depending on HR gene-defect status both at mutational as well as signalling activity level, in particular hypoxia, demonstrating physiological deviations which may be reliant on BRCA defects specifically.

### Developing a transcriptional signature of BRCA1/2 deficiency

To further explore the functional consequences of HRD, we sought to develop a transcriptional signature reflecting the gene expression profiles of HRD and HR-proficient breast cancers. We aimed to ensure that the signature encompassed the various forms of HRD that could be driven by different factors, such as BRCA1 or BRCA2 defects as well as BRCA-independent mechanisms (HRD-BRCA+). To this end, we trained a multinomial elastic net regression model on the expression profiles from two thirds of the TCGA-BRCA cohort to distinguish between different forms of HRD or HR proficiency, with the remaining TCGA-BRCA samples used for testing (Fig. 3a, Methods). A multinomial elastic net approach was chosen due to the ability to remove uninformative features, whilst also tolerating correlated variables. Additionally, since we were seeking a cancer cell-specific phenotype, as opposed to a signal of the tumour microenvironment which can confound bulk sequenced samples, we conducted expression deconvolution using BayesPrism(69) on the training cohort and used the estimated cancer-specific transcriptional profiles for signature development (see Methods). The estimated cell type fractions from BayesPrism significantly correlated with tumour purity estimates(70), indicating reliable cancer cell-specific estimation (Supplementary Figure 13).

**Figure 3.**
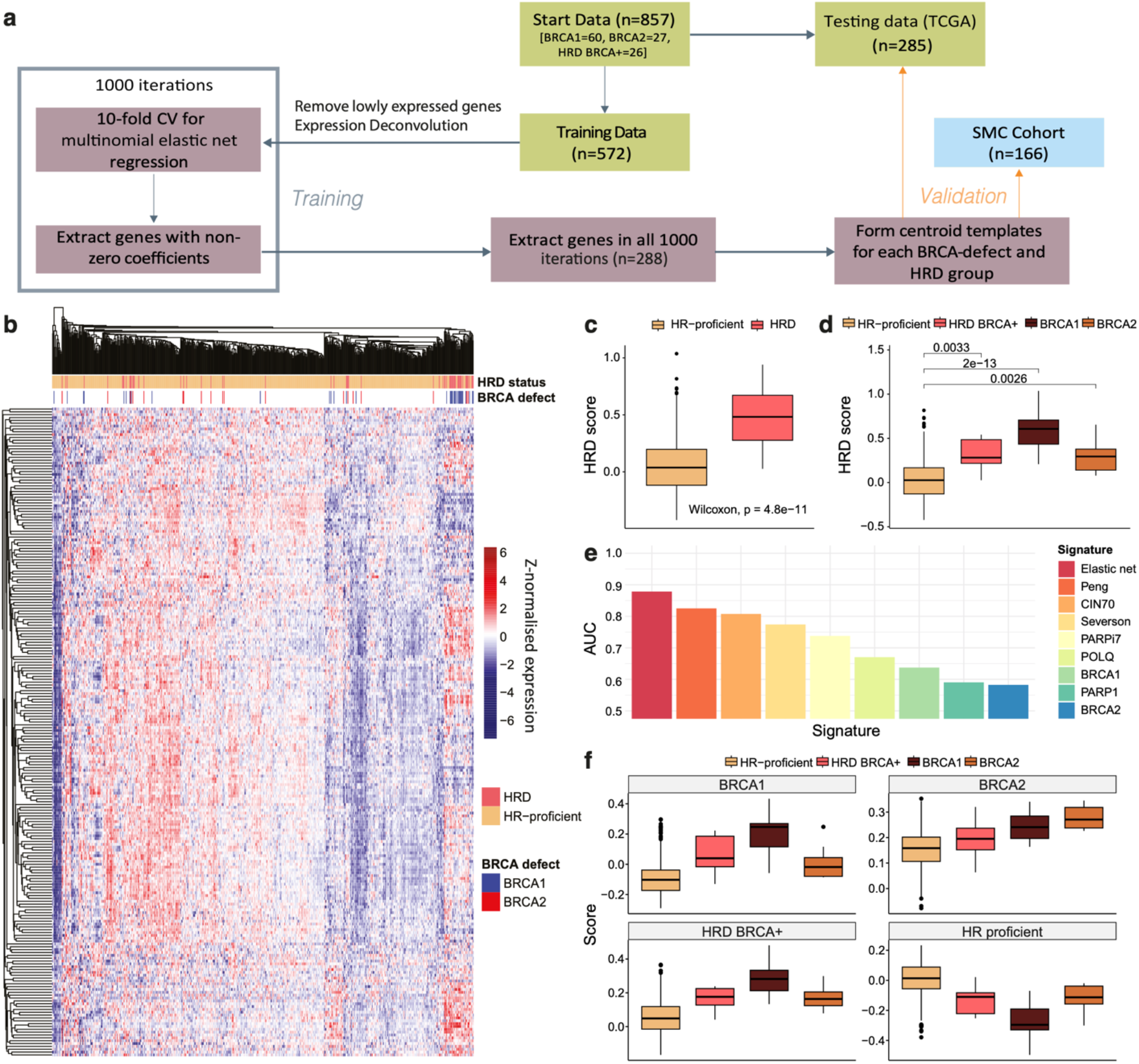
Development and validation of a BRCA defect type-specific HRD transcriptional signature. (a) Workflow for transcriptional signature development. Data is split into training and testing cohorts. The training data undergoes expression deconvolution to extract a cancer cell-specific signal, using the Qian et al. single-cell RNA-seq cohort as a reference, and genes that are lowly expressed in this dataset are removed. Processed training data undergoes 1000 iterations of 10-fold cross validation of elastic net regression, and a signature is formed from the 228 genes selected in every iteration. Centroid templates are formed for HRD/HR-proficient and BRCA-type HRD groups from the 228 genes across the training cohort, and scores for testing and validation cohorts are calculated by correlating the new sample against each template. (b) Summary of the 228-gene HRD transcriptional signature profiles across the TCGA training set. The HRD status assignment is annotated along with BRCA1/2 defects. (c-d) Comparison of HRD scores calculated using the transcriptional signature between c) HRD vs HR-proficient and d) HRD/BRCA-defect groups. (e) Comparison of HRD transcriptional signatures and gene expression markers for predicting HRD status in the TCGA training set, measured by AUC. ‘Elastic net’ refers to the 228-gene transcriptional signature presented in this study. ‘Peng’, ‘CIN70’, ‘Severson’, and ‘PARPi7’ refer to alternative transcriptional signatures as described in the Methods. POLQ, BRCA1, PARP1, and BRCA2 are gene expression markers. (f) Comparison of HRD/BRCA-defect scores across HRD/BRCA-defect groups in the TCGA testing cohort. Each panel corresponds to a specific HRD/BRCA-defect signature, with the y-axis representing correlation with the respective centroid model. Each box represents refers to the samples within the respective group.

The transcriptional signature was generated by conducting 1,000 iterations of multinomial elastic net regression with 10-fold cross validation and extracting the genes which were included in all 1,000 iterations (see Methods). An HRD ‘score’, as well as four scores representing HR-proficiency, BRCA1ness, BRCA2ness, and HRD-BRCA-positivity, were then calculated using a centroid-based approach (see Methods). This procedure was conducted with varying regularisation parameters, as well as with and without observation weights to account for imbalanced groups (see Methods). The final signature, containing 228 genes (Fig. 3b, Supplementary Table 4), was selected on account of its optimal capture in single cell data based on the Qian et al. cohort(71) (Supplementary Figure 14).

The resulting HRD score was adept at classifying samples labelled as HRD or HR-proficient according to the mutational signature-based classifier (AUC = 0.88) (Fig. 3c). An advantage of the signature was that whilst BRCA1-defective samples showed the highest HRD scores, BRCA2-defective samples also demonstrated HRD scores greater than HR-proficient samples, demonstrating that this signature can capture overall features of heterogeneous HRD (Fig. 3d). For HRD/HR-proficiency classification, the signature outperformed other transcriptional signatures associated with HRD(32), BRCA1ness(31), chromosomal instability (CIN70)(34), and PARP inhibitor sensitivity (PARPi7)(33), as well as gene markers associated with HRD (Fig. 3e).

The signature also showed potential at classifying samples depending on their specific BRCA-defect/HRD status, including when applied to BRCA2-defective samples (Fig. 3f; Supplementary Figure 15). The elastic net signature outperformed all others in characterising BRCA1ness, BRCA2ness and HR-proficiency, further demonstrating that, unlike alternative HRD classifiers, using a multinomial approach prevents the resulting signature from skewing away from BRCA2ness, ensuring that the established HRD heterogeneity is captured. Whilst the overall distribution of HRD scores is greater for ER-negative samples in the TCGA testing data regardless of HRD/HR-proficiency status, HRD is also identifiable using the signature after accounting for breast cancer subtype, suggesting that it is not just merely capturing the ER status of the tumours (Supplementary Figure 16).

These results were validated in the SMC breast cancer cohort(42). The 228-gene signature outperformed alternative methods in HRD classification (AUC = 0.81) and demonstrated adept capacity for BRCA-defect and HRD BRCA+ classification in comparison with alternative methods (Supplementary Figure 17). It is noted that HRD classification capacity is slightly reduced in the SMC cohort which is suspected to be due to the small number of BRCA1-defective patients in the SMC cohort, whilst BRCA1-defective patients dominated the HRD group in TCGA.

According to gene set enrichment analysis, DNA repair processes are dominantly enriched across the signature, as driven by *BRCA1*, *BRCA2*, *TOP3B*, and *FANCI* (Supplementary Figure 18). Intriguingly, the signature was also enriched for genes associated with insulin signalling and glucose transport (*IRS1*, *IRS2*, *CACNA1D*, *SOCS3*, *PRKCZ*), and autophagy and mTOR signalling (*RPTOR*, *RRAGD*, *GABARAP*, *ATP6V1E2*, *ATP6V1C1*).

### Key transcriptional contributors to HRD classification

Whilst the 228-gene signature predicts both the HRD and BRCA defect status, we were interested to explore whether a reduced signature could characterise HRD sufficiently. To achieve this, we employed graph attention networks (GATs) that would help us prioritise genes in the signature which have the greatest contribution to distinguishing HRD and HR-proficient phenotypes (Fig. 4a, Methods). The model makes use of the original genes from the signature and the degree to which they are correlated in their expression within the TCGA-BRCA cohort to classify patients as HRD and or HR-proficient, while the correlated gradient information decides which gene sub-modules might drive the classification. Briefly, a patient-specific weighted gene co-expression graph is extracted using weighted correlation network analysis (WGCNA)(72) and these graphs are input into the GAT, which is then trained to distinguish HRD from HR-proficient samples and then selects parts of the graph based on its gradients. The resulting graph neural network showed high accuracy for HRD classification (AUC = 0.90). By ranking the genes using a gradient-based approach for their performance in classifying HRD or HR-proficiency, we were then able to highlight the key genes driving correct model prediction.

**Figure 4.**
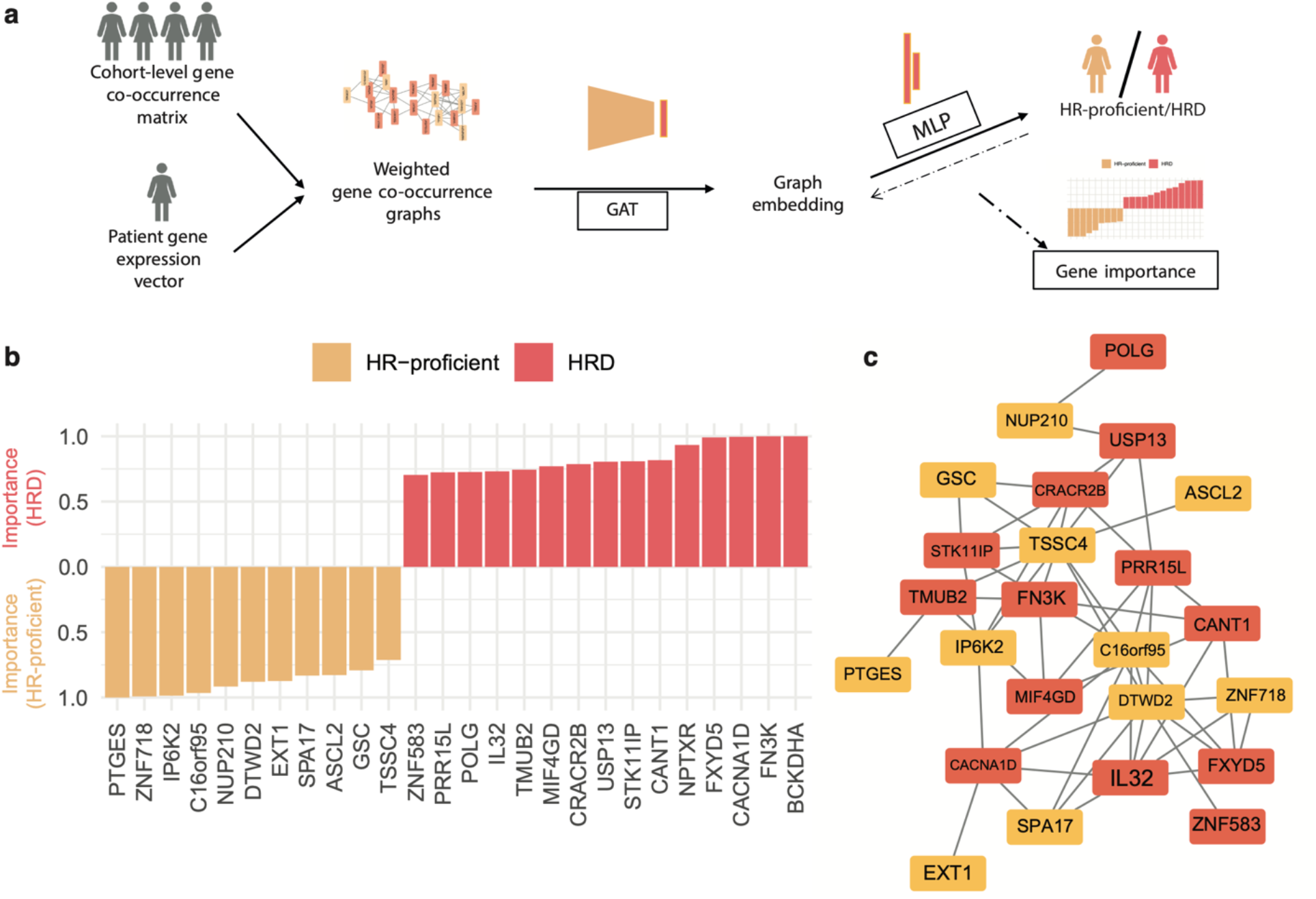
Graph analysis to determine transcriptional signature drivers. (a) Workflow for graph attention network analysis to classify HRD/HR-proficient TCGA-BRCA patients and determine gene importance. Weighted gene co-expression graphs are built from the gene expression profiles of the TCGA-BRCA cohort, while taking into account the patient-level gene expression for the 228 genes in the transcriptional HRD signature. A graph attention network (GAT) is then trained to distinguish HRD and HR-proficient samples using the weighted co-expression graphs as inputs. The output highlights part of the graphs with greater weight in the classification and generates an importance score for each genes. (b) The top-ranked 26 genes in the HRD versus HR-proficiency classification with importance scores greater than 0.7. The colour indicates the phenotype (HRD/HR-proficient) for which the gene is predictive. (c) The co-expression graph of 24 of the 26 highly ranked genes for classification. Genes are connected only if they are co-expressed in the cohort, and genes with no connections have been removed. The colour of the nodes depicts the associated phenotype as in (b).

The analysis highlighted 26 genes which were sufficiently important for classifying the HRD and HR-proficiency groups, of which 15 were predictive of HRD and 11 were predictive of HR-proficiency (Fig. 4b-c, Supplementary Table 5). A number of these genes have been associated with DDR, including *USP13*, a regulator of replication stress involved in ATR activation via TopBP1 deubiquitination(73,74), *POLG*, a key regulator of mitochondrial DNA replication and repair(75,76), and *IP6K2*, a stabiliser of DNA-PKcs and ATM leading to p53 phosphorylation(77,78). Additionally, the reduced signature contains two genes encoding zinc finger proteins (*ZNF718* and *ZNF583*) which are involved in both HR and NHEJ(79,80). On their own, these 26 genes provide an adept reduced gene signature for capturing HRD, with maintained capacity for HRD classification across BRCA2-defective samples (Supplementary Figure 19).

### Testing the signature against sensitivity to PARP inhibitors

To determine the therapeutic relevance of the transcriptional signature of HRD, we next applied the signature to breast cancer cell lines to predict PARP inhibitor sensitivity. The signature was applied to 68 breast cancer cell lines from the Cancer Cell Line Encyclopaedia (CCLE)(81), and matched to PRISM drug sensitivity data from 26 of these cell lines (Supplementary Table 6).

Cell lines with increased transcriptional scores for HRD showed increased sensitivity to four different PARP inhibitors, as represented by lower PRISM scores (Fig. 5a). We note that the correlation is marginal, especially regarding olaparib and talazoparib. However, the 228-gene signature still shows a stronger correlation with PARPi sensitivity in these cell lines than any alternative signatures (Supplementary Figure 20). This is likely a result of the signature being developed from bulk-sequenced primary tumour samples, whilst the cell lines will be lacking a microenvironmental component. Whilst we have attempted to account for microenvironmental signals using expression deconvolution, these extracted cancer cell-specific signals will still exist in a broader environmental context which the cell lines will be lacking, hence a resulting transcriptional HRD signal will likely vary significantly.

**Figure 5.**
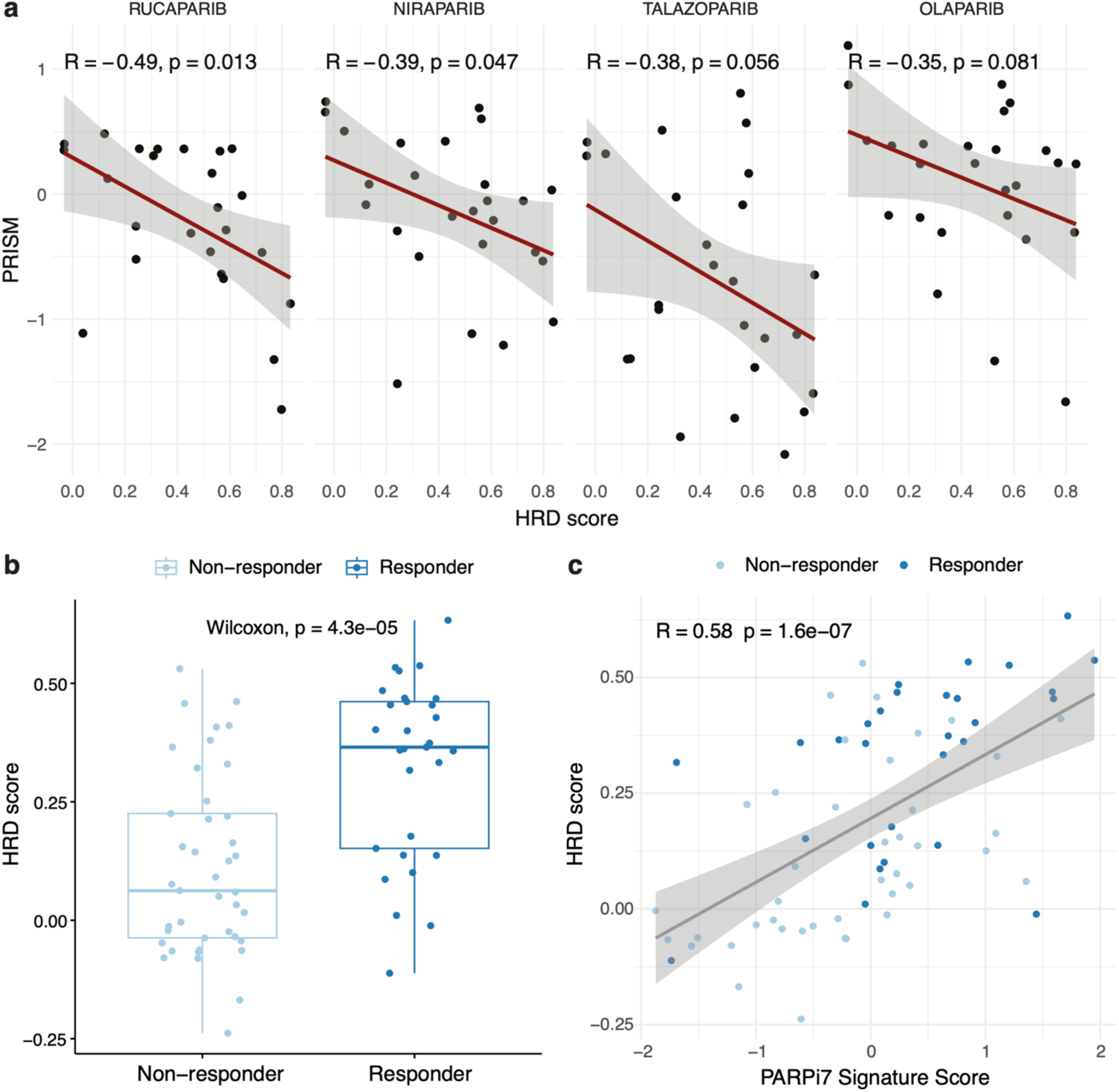
HRD transcriptional signature is linked with PARP inhibitor sensitivity in breast cancer cell lines and patients. (a) Correlation between the HRD transcriptional scores calculated using transcriptomes from breast cancer cell lines from CCLE and sensitivity to PARP inhibitors evaluated using the PRISM metric. (b) Comparison of HRD transcriptional scores between responders and non-responders to olaparib and durvalumab combination treatment in the I-SPY2 trial. (c) Correlation of HRD transcriptional scores against the PARPi7 signature score calculated by Pusztai et al.(38) in the I-SPY2 treatment arm patients. Responders are more frequently scoring high using our signature compared to PARPi7.

The HRD score also predicted responses to combined PARP and checkpoint inhibition in breast cancer patients. The signature was applied to 105 HER2-negative, Stage II/III breast cancer patients from the I-SPY2 trial(38). In this trial, 71 patients were treated with a combination of olaparib and the PD-L1 inhibitor durvalumab, as well as the neoadjuvant chemotherapy taxol, whilst 34 control patients were treated with taxol alone. Amongst the patients in the treatment arm, 29 showed pathologic complete response (pCR), and these patients showed significantly increased HRD scores compared to those who did not display pCR (Fig. 5b). Our HRD score was significantly correlated with their own PARPi7 signature score from the trial but showed a better separation between responders and non-responders (Fig. 5c, Supplementary Figure 21). Overall, this demonstrates that, in capturing a general transcriptional phenotype of HRD, the 228-gene HRD signature is also linked with PARP inhibitor sensitivity, especially in patient samples.

### Application of the HRD signature to single cells

When considering HRD in the context of treatment or identification, it is often forgotten that whilst HRD can manifest and cause effects at the level of the whole tumour, it is still an intrinsically cellular phenotype. Since the transcriptional signature was developed with the intention of characterising HRD in terms of tumour-specific signalling, as opposed to that of the microenvironment, this suggested that the signature could be applied to scRNA-seq data, which would provide insight into the distribution of HRD across cells and generate further questions about the varying roles of HRD and HR-proficient tumour cells within the context of the tumour microenvironment.

To investigate whether the signature may be applicable to single cell data, we first applied it to 11 breast cancer samples with matched bulk and single cell-sequencing data from Chung et al(82). We show a good correlation between the bulk HRD scores and the mean HRD scores across the individual tumour cells within each sample (Fig. 6a), providing an indication that the HRD signal captured in bulk sequencing data reflects, on average, the levels seen in single cells.

**Figure 6.**
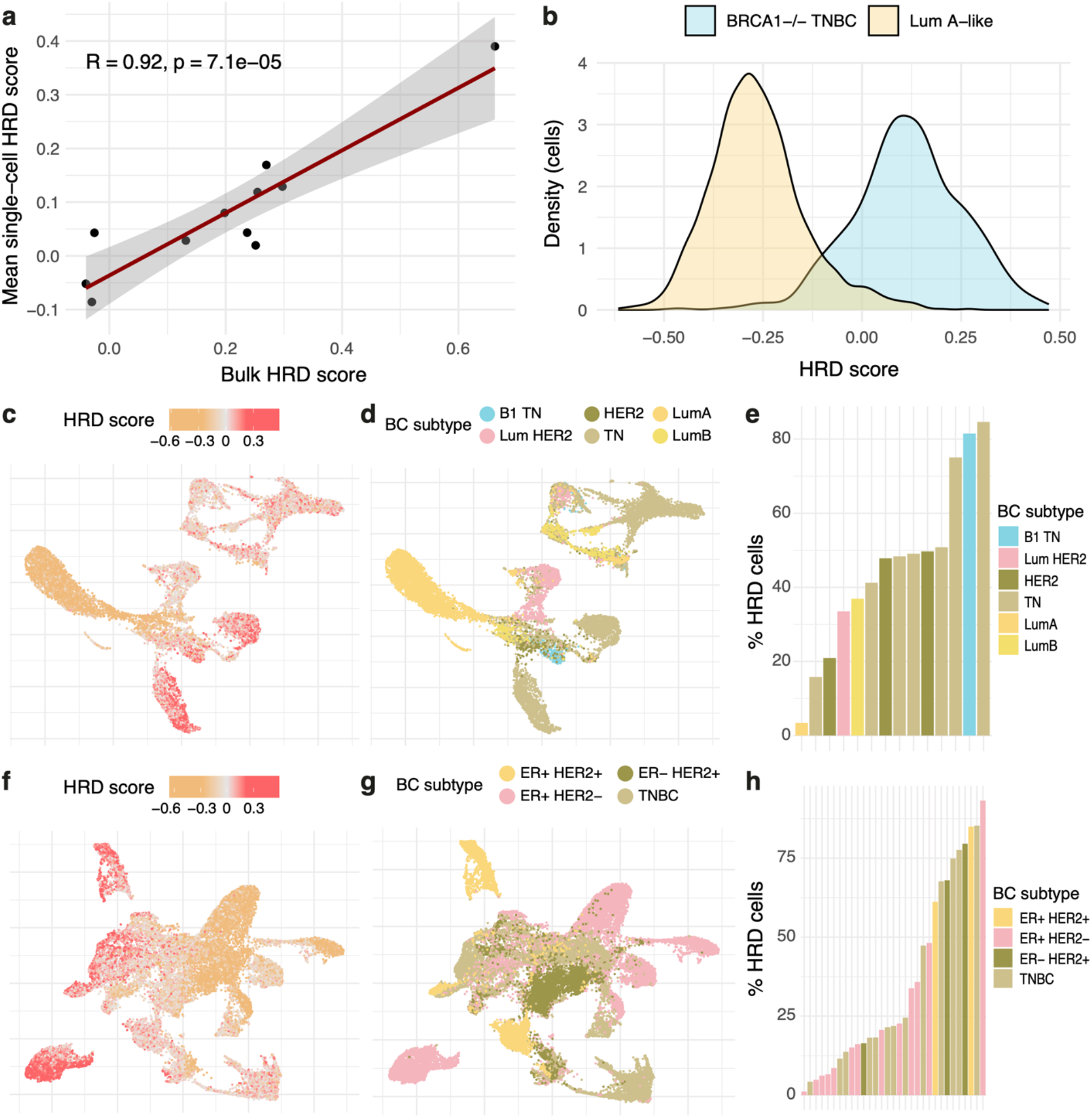
Transcriptional profiling of HRD in single cell sequenced breast cancer cells. (a) Correlation of mean HRD transcriptional score across individual cancer cells against matched bulk RNA sequencing from the Chung et al cohort(108). (b) Distribution of HRD scores across tumour cells from a Stage III BRCA1-defective TNBC sample (sc5rJUQ033) and a Stage II Luminal A sample (sc5rJUQ064) from the Qian et al cohort(71). (c-e) Profiling of HRD across tumour cells from the Qian et al cohort as demonstrated by UMAP coordinates labelled by c) HRD score and d) breast cancer subtype. (e) The proportion of cells within each sample with HRD scores greater than zero in the Qian et al cohort. The defined breast cancer subtypes include here are: ‘B1 TN’ = BRCA1-defective Triple negative, ‘Lum HER2’ = Luminal-HER2+, ‘HER2’ = HER2 positive, ‘TN’ = Triple negative, ‘LumA’ = Luminal A-like, ‘LumB’ = Luminal B-like (f-h) Profiling of HRD across tumour cells from the Bassez et al. cohort(85), similar to (c-e).

Following this, we applied the signature to a cohort of 14 single cell-sequenced breast cancers containing over 44,000 cells from Qian et al(71). These 14 samples included one BRCA1-defective sample (sc5rJUQ033), which we assumed to be HRD, and one Stage II Luminal A sample (sc5rJUQ064) which we assumed to be HR-proficient, given the characterisation of this type of breast cancer as slow-proliferating(83,84). These two samples display substantially different distributions of HRD scores across the cancer cells in these respective samples, with the Stage II Luminal A sample displaying a distinctively more HR-proficient distribution (Fig. 6b), further demonstrating that a transcriptional signature of HRD can potentially be captured at single-cell resolution.

Across the Qian et al. cohort and 31 treatment-naïve samples obtained from the Bassez et al. cohort(85), on average 10.8 and 11.2 genes from the signature were expressed per cancer cell, respectively (Supplementary Figure 22a-b). The mean number of genes in the signature expressed per cell across each samples varied between 6.74 and 17.9 in the Qian et al. cohort (Supplementary Figure 22c) and 6.40 and 23.3 in the Bassez et al. cohort (Supplementary Figure 22d), indicating sufficient capture of the transcriptional signature across the single-cell cohorts.

The cancer cells from these 14 samples displayed intra-sample heterogeneity of HRD scores (Fig. 6c; Supplementary Figure 23a) that matched the clustering by breast cancer subtype (Fig. 6d). Generally, the triple negative and BRCA1-defective samples presented a greater proportion of HRD cells, defined as displaying a transcriptional score greater than zero (Fig. 6e).

Similarly to Qian et al(71), the tumour cells obtained from Bassez et al.(85) displayed a distinctive gradient of HRD scores (Fig. 6f; Supplementary Figure 23b) in accordance with the clustering by breast cancer subtype (Fig. 6g), and TNBC samples displayed greater proportions of HRD cells in comparison with receptor-positive samples(Fig. 6h). Moreover, we found that, whilst the HRD scores from the TME were consistent across samples and tended to centre closely around zero, these scores varied far more broadly across samples within the cancer cells across both cohorts, indicating that the transcriptional signal of HRD is likely arising strongly from the tumour cells (Supplementary Figure 23c-d).

This further demonstrates the potential provided by this transcriptional signature of HRD to capture this phenotype at single-cell resolution, as well as demonstrating the heterogeneity of HRD levels across individual samples.

### Exploration of the HRD tumour microenvironment at single cell level

To further explore the activities of HRD and HR-proficient cells within the tumour microenvironment, we applied CellphoneDB(86), an extensive database of ligand-receptor interactions, to observe whether the interactions established between tumour cells and the surrounding immune and stromal cells varied based on HR capacity utilising both the Qian et al and Bassez et al cohorts(71). In both cohorts, cell-cell interactivity profiles were dominated by fibroblast, endothelial, myeloid, and dendritic cells, with cancer cells demonstrating fewer interactions with the TME (Fig. 7a-b). However, across all cell types, HRD cancer cells consistently displayed fewer significant interactions with the TME, in particular as the target of these interactions, than HR-proficient cells (Fig. 7c-d).

**Figure 7.**
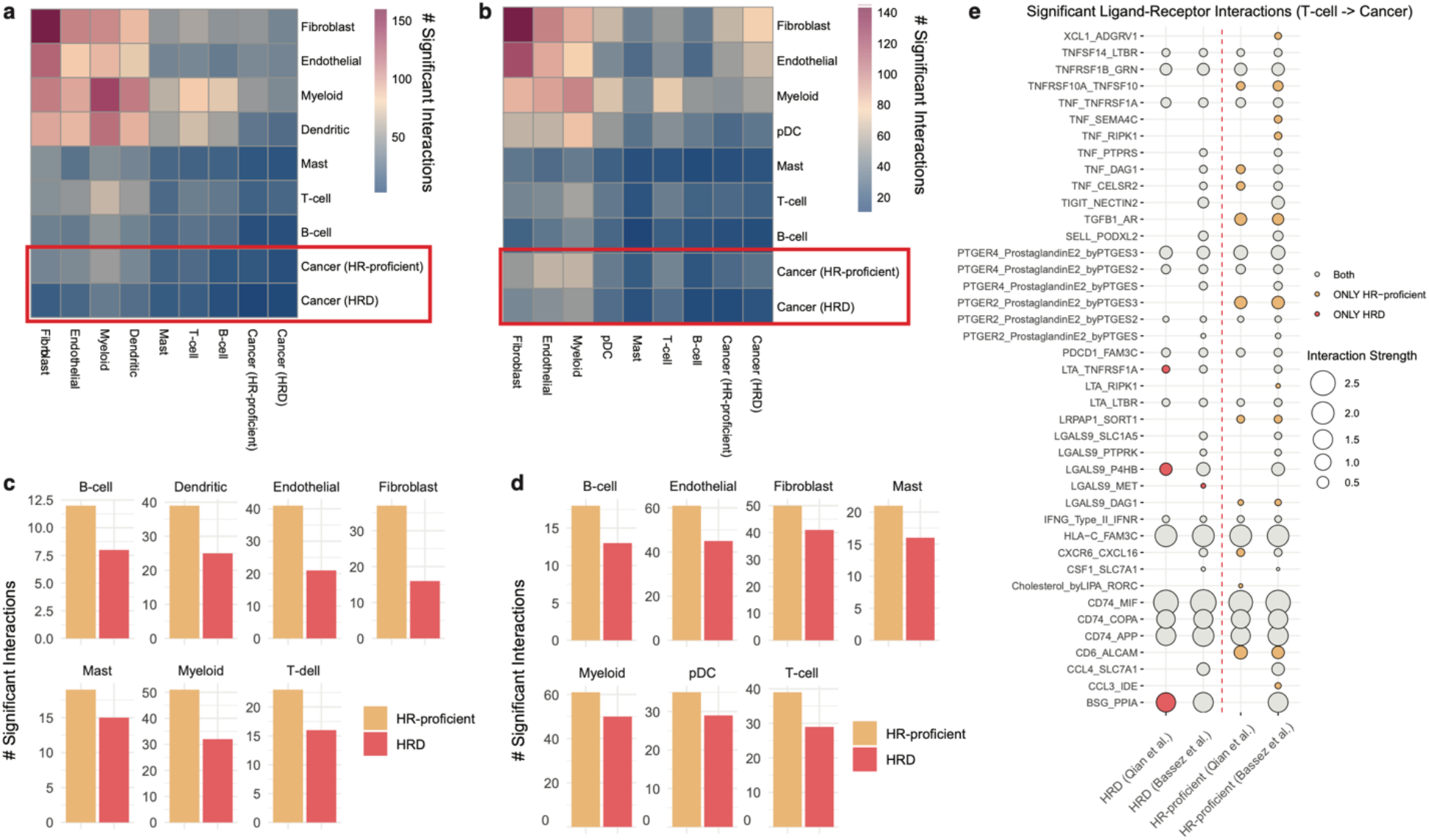
TME-cancer interactivity across HRD and HR-proficient cancer cells. (a-b) Number of significant ligand-receptor interactions established between cells in the (a) Qian et al.(71) and (b) Bassez et al.(85) cohorts according to CellphoneDB. Cancer cells are labelled as HRD if they have a positive HRD score, HR-proficient otherwise. The x-axis refers to cell types as sources, and the y-axis refers to cell types as targets. (c-d) The number of significant interactions between TME cell types as sources and cancer cells as targets, separated by HR status, across the (c) Qian et al. and (d) Bassez et al. cohorts. (e) Specific ligand-receptor interactions between T-cells and cancer cells, with cancer cells as the targets, across the Qian et al. and Bassez et al. cohorts. The red circles indicate interactions unique to HRD cells within a given cohort, and the yellow circles indicate interactions unique to HR-proficient cells within a given cohort. Grey circles represent common interactions.

While some common interactions were observed, multiple ligand-receptor pairs uniquely mediated the interactions of HR-proficient cells with T cells across both cohorts (Fig. 7e). In particular, these included interactions involving TNF, TGFb, and prostaglandin E2 signalling across both cohorts and from different cell types (Supplementary Figure 24). These results align with a previous hypothesis suggesting downregulation of TNF signalling as a mechanism of cell survival following BRCA2 deficiency on account of the resulting decrease in caspase-induced apoptosis(55). Unique TME-HRD cell interaction pairs often involved *LGALS9* signalling, a feature which also re-occurred across cell types and cohorts (Supplementary Figure 24). *LGALS9* has previously been associated with antimetastatic potential in breast cancer(87), potentially indicating a compensation for increased chromosomal instabilility which may drive metastasis(88). Additionally, expression of the *LGALS9* gene product, Gal-9, is increased following taxane treatment in TNBC, due to nuclear activation of NF-kB, which is also upregulated in HRD breast cancers(89,90).

This analysis unveils the specific strategies HRD and HR-proficient cells employ in their crosstalk with the TME, highlighting differences in the type and variety of molecular components involved. Generally, the HRD cells appear less “promiscuous” while HR-proficient cells display a wider array of interaction strategies. However, these findings do not inform us on whether HRD cells respond less or more frequently to other cells in their environment, as the individual cell-cell interactions cannot be inferred using this method, and should not be interpreted as such.

Overall, our results highlight a complex pattern of interactions between cancer and non-cancer cells which may be mediated by the DNA damage response and could, in the long term, inform treatment strategies that jointly target HRD tumours and their microenvironments.

## DISCUSSION

As the clinical utility and capabilities of personalised treatments increase, it is necessary to ensure that the features predicting positive treatment response can be identified reliably. In this study, we present multi-scale approaches to characterising HRD which can be applied in a variety of contexts. We developed a method for high-confidence HRD identification in exome sequenced breast cancers which incorporates indel events that indicate both the presence of HRD and alternative DSB repair mechanisms employed in the event of HRD. We demonstrate that even small amounts of indels like the ones expected to be seen in WES data improve HRD classification. Furthermore, the HRD group defined by our genomic signatures displays the characteristic features expected of such cancers, including *MYC* amplification and elevated *POLQ* expression. Applying this classifier to the TCGA-BRCA cohort, we then developed a 228-gene transcriptional signature that characterises the heterogeneity of HRD, whilst also correlating with PARP inhibitor sensitivity and displaying the capacity to define HRD at single cell resolution.

In creating the mutational signature-based classifier, we only considered SBS and indel signatures due to our focus on the HRD phenotype and data availability in TCGA. However, the generalisability of the method is worth highlighting. Copy number signatures could be effectively integrated into this method to improve HRD classification, especially if copy number profiles can be divided into specific features and contexts as demonstrated in previous studies(20,21). Furthermore, our method also defines subgroups enriched for alternative mutational processes, including APOBEC cytidine deamination and mismatch repair deficiency. With regards to this, double base substitutions (DBSs) can also be included within the method and may be of use for improved classification of processes with associated DBS signatures, such as mismatch repair deficiencies and tobacco-associated mutagenesis.

We applied a probability threshold of 0.79 for HRD classification using the mutation-based classifier, which was determined through optimising the resulting F-score for identifying patients with HR gene defects. Whilst using this threshold leads to high-confidence classification of HRD in samples which do not harbour HR gene defects (Supplementary Figure 11b), this does lead to a minor but notable decrease in sensitivity. Additionally, we note that these groups may not be distributed identically in the TCGA cohort, as demonstrated by the differences in rates of BRCA defects (13.6% in ICGC compared to 8.99% in TCGA), suggesting that the prior distributions, whilst based on real data, might not be wholly representative. It is non-trivial whether to emphasize sensitivity or specificity when generating an HRD classifier, in which it is known that the specificity should not be 100%. This is because a key difficulty in developing an HRD classifier is the lack of ground truth beyond HR gene defects and, as was used for SigMA and the simulation analysis conducted in this study, SBS3 signature contribution in WGS data. Therefore, we used a balanced F-score which considers both with equal importance. However, we note that different probability thresholds can be applied, as was utilised for SigMA(27).

Mutational signatures have also shown promise for HRD classification in targeted panel sequencing, when presented as a likelihood-based approach as was done through SigMA(27). Due to the availability of gene panels, this development presents invaluable clinical relevance. Whilst we also employ a likelihood-based approach, owing to the substantially decreased indel loads identified through targeted panel sequencing it is unlikely that this method would significantly contribute to improved HRD classification, and therefore do not recommend its application to gene panels. Furthermore, it should be noted that our exome classifier’s specific clinical utility is limited. Whole genome sequencing will likely become increasingly available for mutational signature-based diagnostics such as HRDetect, and panel sequencing is already widely applied for identifying gene defects. However, the primary benefit of an exome-based classifier is its application to large-scale genomics resources such as TCGA, enabling further profiling of HRD from a broad range of omics perspectives and new hypotheses generation which these resources enable.

Whilst it is not surprising that our 228-gene transcriptional signature outperforms alternative methods given that it was trained using labels determined by our own mutational HRD classifier, this transcriptional signature also simultaneously captures BRCA1- and BRCA2-specific deficiency phenotypes, highlighting the distinct consequences of loss of function of these two genes. It is worth noting that we additionally capture a group of tumours that display transcriptional profiles closer to those of classically HR-deficient BRCA mutated samples but lacking any BRCA defects. These tumours might be experiencing some level of HRD-like state due to more complex changes across the HR and linked pathways, some of which may be epigenetic or of other nature. Further analyses are needed to shed light into the aetiology of these cancers, and it is likely they are a rather heterogeneous group.

In terms of its relevance in a therapeutic context, we found that our HRD transcriptional signature was more strongly associated with PARP inhibitor sensitivity in patients than in cell lines. Given that the signature was developed using breast cancer patient samples, this was likely to have been the case. While we have attempted to ensure that we are capturing a tumour-cell intrinsic HRD signature by correcting for microenvironmental signals in bulk data, some tumour intrinsic regulation might still be partly environmentally triggered, and this component would not be captured in cell lines which lack this microenvironment. Additionally, due to a broad variety of factors including genetic instability and growth conditions, drug responses across cell lines may be hugely variable(91), which may partially explain the decrease in signature performance when applied to cell lines.

Future developments are likely to focus significantly on PARP inhibitor resistance. Archetypal mechanisms of PARP inhibitor resistance are becoming well established, such as BRCA1 reversion cases, 53BP1 loss following BRCA1 loss, and PARG loss following BRCA2 mutations(92), and features such as reprogramming of cell survival pathways and an increasingly mesenchymal phenotype have been associated with gradual PARP inhibitor resistance(93). Currently there are no available datasets demonstrating the effect of HR resurgence on tumour heterogeneity, however these will provide an invaluable resource for studying HRD moving forward.

Additionally, recent developments in inferring mutagenic processes in single cells have shed light on the driving forces behind the evolution of tumour heterogeneity in TNBC and high grade serous ovarian carcinoma (94). In breast cancer, HRD tumours are generally believed to be more immunogenic due to their potential to generate increased mutational loads and neoantigen signalling, which can be exploited for checkpoint inhibition(43,95,96) BRCA defects have also been associated with increased immunosurveillance in high grade serous ovarian cancer, and CellphoneDB was recently applied to highlight a malignant cell population associated with poor prognosis in ovarian cancer, which was associated with immune cell interactions and displayed generally low levels of chromosomal instability(97,98). Additionally, CellphoneDB was used to explore human breast cancer immune microenvironments, and specifically highlighted a large number of unique interactions between fibroblasts and endothelial cells, as well as smaller levels of interaction by T-cells and B-cells, as we have also observed, which was attributed to fewer genes being expressed in these cell types(99). While the TME has been fairly extensively explored in breast cancer bulk datasets and more recently in single cells, our understanding of how the tumour-TME crosstalk is established in the context of HRD or HR proficiency at single cell resolution is much more limited. We show that our transcriptional signature may be employed to highlight patterns of HRD in single cells, and this paves the way for further explorations into the way that DNA repair deficiencies may influence their microenvironments, and vice versa. Thus, we believe our analysis can pave the way to more detailed interaction studies by highlighting specific strategies of tumour-T cell coupling in HRD cells which may have relevance to immunotherapy effectiveness. The association between HRD and various facets of the tumour microenvironment is already being mined for therapeutic potential through investigation of joint PARP and immune checkpoint inhibition(38,100,101), and chromosomal instability has been shown to elicit an inflammatory response offering further targets for combination treatment(55,102). The capacity to investigate differential cell-cell interactivity of HRD and HR-proficient cells may allow for insight into further mechanisms which may be exploited.

## CONCLUSIONS

We have demonstrated that HRD classification in exome sequenced breast cancers can be improved by leveraging the presence of HRD-associated indel events, and have shown that mutational and phenotypic profiles of HRD persist regardless of the presence of HR gene defects. These classifications have been used to develop a transcriptional signature which is associated with sensitivity to PARP inhibitors and can be applied to characterise HRD in single-cell RNA-sequencing data. In examining the TME crosstalk at single cell resolution, we demonstrate substantial variation in cell-cell interactivity patterns dictated by the HRD or HR proficiency status of the tumour cells, suggesting distinct pathways mediating immune recognition and/or escape. These findings pave the way to further investigation of the heterogeneity of HRD and HR-proficiency both at patient and individual cell level, as well as therapeutic implications.

## MATERIALS AND METHODS

### Data sources

Whole-genome sequencing data from the ICGC-BRCA cohort was obtained from DCC data release 28(103). Data from two projects was included: BRCA-UK (n = 45) and BRCA-EU (n = 569), resulting in a total of 614 samples. Exome sequencing data from TCGA was obtained from the GDC Data Portal using the TCGAbiolinks R package(104). Somatic mutation data collated using the Mutect2 pipeline was obtained using the GDCquery() function. Annotation of BRCA-defective samples in TCGA was taken from Valieris et al.(105). Samples were considered BRCA1/2-, RAD51C-, or PALB2-defective if they were assigned either ‘Bi-allelic inactivation’ or ‘Epigenetic silencing’. HRD index scores as calculated by Myriad for TCGA samples were obtained directly from Marquard et al.(22). CX3 scores were obtained from Drews et al.(20).

Exome sequencing and transcriptional profiling from the SMC cohort of Korean breast cancer patients, as well as chromosome arm-level copy number alterations for the TCGA-BRCA cohort, were downloaded from cBioPortal(106).

FPKM-normalised gene expression data for TCGA was obtained from the GDC Data Portal. The proliferation/cell cycle arrest capacity of the tumours was calculated from RNA-seq profiles using the quiescence (Q) score defined in Wiecek et al. (107), and was defined as 1-Q, with positive scores indicating a higher proliferative capacity.

Cell line expression data and PARP inhibitor sensitivity profiles were obtained from the Cancer Cell Line Encyclopaedia (CCLE)(81). Transcriptional profiling and clinical data of patients from the I-SPY2 trial were collected from the Gene Expression Omnibus (GEO) database, with accession number GSE173839.

Single-cell RNA-seq data from Chung et al.(108) was collected from GEO database through the accession numbers GSE75688. Treatment-naïve single-cell RNA-seq data from Qian et al.(71) and Bassez et al.(85) were downloaded directly from the Lambrechts laboratory website https://lambrechtslab.sites.vib.be/en/team. Single-cell RNA-seq profiling from Qian et al. and Bassez et al. were processed using the Seurat R package(109), to extract only cancer cells with between 200 and 6,000 unique feature counts and mitochondrial content less than 15%, the expression profiles of which were then log-normalised.

### Generation of HRD classifier for exome sequenced breast cancer samples

#### Mutational signature contributions in WGS breast cancer samples

Mutational signature analysis of 614 WGS breast cancer samples obtained from the International Cancer Genome Consortium (ICGC) project was conducted using the deconstructSigs R package(40). SBS and indel signatures from the COSMIC v3.3 database were included if they appeared in >1% breast cancer samples according to the Pan Cancer Analysis of Whole Genomes (PCAWG) project(17). SBS and indel contributions were calculated separately, and the results were combined for subsequent clustering analysis.

#### Formation of HRD-specific mutational spectra

The 614 ICGC breast cancer samples were clustered according to their SBS and indel signature contributions, which were calculated separately using the deconstructSigs R package(40). Since we intended to use WGS samples to calculate estimated signature contributions within the exome regions, we normalised the mutation profiles to account for the frequency of each mutation type occurring in the exome relative to the whole genome. For SBS profiles, this was done by setting the option tri.counts.method = ‘genome2exome’ in deconstructSigs. For indel profiles, we used the ICGC breast cancer cohort to count the total frequency of each of the 83 indel mutation types, and estimated the frequency of each within the exome by excluding mutations appearing within intergenic regions. The indel profiles of each sample were then multiplied by the ratio of the frequency of each mutation type within the exome to the whole genome. Additionally, mutation spectra in exomes may also differ from the rest of the genome on account of transcription-coupled damage and repair, which cannot be accounted for in WGS samples even after factoring for varying triplet frequencies.

Model-based clustering was conducted using the mclust R package using finite mixture modelling(110). The final classification and optimal number of clusters was selected according to the Bayesian Information Criterion value. Whilst 22 clusters were initially identified via this method, two of these clusters consisted of only one sample each, neither of which displayed discernible features, therefore these clusters were discarded. The 20 remaining clusters were named according to the most prevalent features, with the seven clusters enriched for SBS3 labelled as ‘HRD’.

The mutational spectrum for each cluster was determined by collectively calculating the mean distribution of the 96 SBS and 83 indel mutation types. The result is 20 representative mutation distributions consisting of mutation events, each summing to one (Supplementary Figure 3).

#### Application of mutational spectra for HRD classification in TCGA

The frequencies of the 179 mutation events across 986 exome-sequenced breast cancer samples obtained from TCGA was determined using the sigminer R package(19). The probability of a sample being assigned to a specific cluster given the set of aberrations displayed follows Bayes’ theorem as follows:

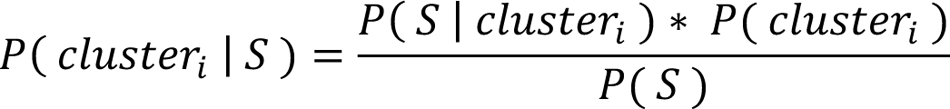

where *S* represents the *n* mutations forming the mutational profile of the sample, *P*(*cluster*_i_) is the prior probability of assignment to cluster *i*, with *i* ∈ [1, 19], as estimated from clustering of WGS ICGC samples, and *P*(*S* | *cluster*_i_) is the likelihood of *S* occurring in a sample from cluster *i*, calculated as:

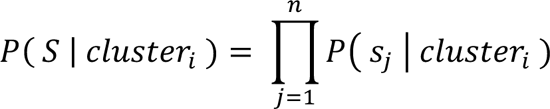

where *S*_j_ is the *j*th mutation, *P*: *S*_j_; *cluster*_j_) is the probability of *S*_j_ within the mean mutational spectrum of *cluster*_i_, and the normalising constant *P*(*S*) is the sum of likelihoods multiplied by prior probabilities for all 19 clusters:

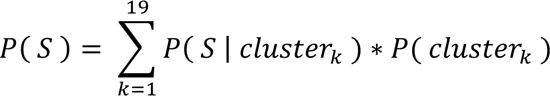

The overall probability of a sample being HRD was calculated as the sum of the probabilities of a sample appearing across the seven HRD clusters, and samples with a probability of greater than 0.79 were deemed HR-deficient. For BRCA defect-specific HRD classification, samples are assigned to the specific cluster to which they have the greatest probability of assignment.

### Evaluation of the HRD classifier

The success of the classifier was determined by calculating its ability to identify patients with BRCA1/2 defects. The characterisation of over 900 TCGA samples for either bi-allelic inactivation or epigenetic silencing of BRCA1 or BRCA2 by Valieris et al.(105) was used as the truth label/gold standard annotation. The F-scores for each HRD classification method were calculated as:

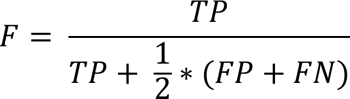

where *TP* is the number of true positives, *FP* is the number of false positives, and *FN* is the number of false negatives. Whilst we expected model sensitivity to be close to 100% (as we assumed that the majority of BRCA-defective samples would be HRD) and did not expect these results for model specificity (due to the known presence of BRCA-positive HRD samples), we aimed to ensure that overclassification of HR-proficient samples as HRD was as limited as possible, and so did not apply a weighted F-score. HRD classification using the HRD index score was done using thresholds of 42 and 63 due to their application in the literature(23,44).

### Simulation Analysis

We tested our method’s HRD classification performance in a low mutation count context using simulated data where we varied the fraction of indels included in a sample. This also allowed us to understand what impact indels generally have for HRD classification. Categories for simulation analysis were generated using hierarchical clustering applied to the SBS signature contributions calculated for the ICGC-BRCA cohort. Samples were downsampled to mutational burdens of 25, 50, and 100, with the additional constraint that the proportion of indels in the simulated data were set from 0 to 0.5, in steps of 0.05. Each combination of sample sizes and indel proportions was iterated 100 times, and differences in the resulting AUC values for identifying SBS3-enriched samples using the classifier at each iteration were analysed using Wilcoxon rank-sum testing.

To analyse the effect of altering the indel contributions within the likelihood distributions, we repeated the analysis except instead of constraining the indel proportions, we multiplied the indel contribution to each likelihood by increasing factors: 1/5, ¼, 1/3, ½, 1, 2, 3, 4, 5.

Simulation analysis was repeated using all 20 clusters by downsampling each sample to sizes of 25, 50, and 100, with no constraint on indel proportions, calculating the AUC for correct cluster assignment using the classifier, and conducting 100 iterations of this process.

### Mutation enrichment analysis

Mutation enrichment analysis was conducted on 738 genes which have been causally implicated in cancer according to COSMIC as of July 3^rd^ 2023 (111). Only genes which were mutated in more than 5% of samples were included in the analysis, and their enrichment in the HRD/HR-proficient groups was calculated using a Fisher’s exact test. Mutated genes under positive selection were identified by dN/dS analysis, which was conducted using the dNdScv R package(54) with default parameters. The analysis was run independently for six groups: all HR-proficient, all HRD, BRCA1-defective, BRCA2-defective, RAD51C-defective, and HRD BRCA+.

### Chromosome arm-level enrichment analysis

For all chromosome arms, enrichment of CNAs in HRD breast cancer samples compared to HR-proficient samples were calculated using Fisher’s exact tests independently testing gains against non-gains (normal or loss), and losses against non-losses (normal or gain).

### Pathway Enrichment Analysis

Differential activity of 14 signalling pathways was analysed using decoupleR(112), which was applied to RNA-seq counts from the TCGA-BRCA cohort. Lowly-expressed genes were removed and the remaining data was VSN-normalised. Differential expression analysis was conducted using the DESeq2 R package(113), and the results of this analysis were fed into the decoupleR R package to estimate pathway activity. Cancer type-specific hypoxia scores, using the Buffa signature(114), were obtained from Bhandari et al.(115).

### Generation of a transcriptional signature of HRD

#### Data preprocessing

Samples from the TCGA-BRCA cohort were selected only if they included exome-sequencing data (and therefore had been assigned as HRD/HR-proficient), and a BRCA defect label obtained from Valieris et al.(105). To prevent confounding, samples harbouring defects in RAD51C or PALB2 were excluded. This resulted in 857 samples, of which two-thirds (n=572) were assigned as training Training and testing sets were defined using the createDataPartition() function from the caret R package(116) to ensure equal proportions of each HRD/BRCA-defect group within the two sets.

#### Expression deconvolution

Expression deconvolution was conducted using the BayesPrism R package, which estimates cell type-specific bulk expression profiles from a single-cell RNA-seq reference dataset. In this case, the Qian et al.(71) dataset was used as a reference dataset. Genes from selected groups, including mitochondrial, ribosomal, and chromosome X and Y genes, were excluded. Following this, protein coding genes only were included. To ensure that the resulting signature would be suitably applicable to single-cell RNA-seq data, genes that were expressed in less than 2% of the cancer cells in the Qian et al. dataset were excluded from further analysis, resulting in 9,853 genes for downstream analysis.

#### Development of the multinomial transcriptional signature

The transcriptional signature was generated by multinomial elastic net regularised logistic regression using the glmnet R package(117). We performed 1,000 iterations of 10-fold cross validation using the cv.glmnet() function with type.multinomial = ‘grouped’. Initially, four signatures were created, setting alpha = 0.25 or 0.5, and applying or excluding weightings to account for group imbalances. We collated the coefficients provided for all features for each iteration in a model generated using λ = lambda.min being the value of λ which gives the lowest mean cross-validated error. The non-weighted elastic net model, with alpha = 0.25, was selected due to its presence within the Qian et al. cohort (Supplementary Figure 14). The final signature was formed of 228 genes which were assigned non-zero coefficients across all 1,000 iterations.

Similarly to Severson et al.(31), the signature was calculated using a nearest centroid method. To create the BRCA1/2-deficiency signature, the TCGA training data was split into its four categories, and a template was created for each group by taking the mean expression of each of the 228 genes across the samples in that category. For new samples, four ‘scores’ were then created by calculating the Pearson’s correlation coefficient between the expression profile of the new sample and each of the four templates.

The same procedure was also applied to generate an HRD signature, except only two templates were created relating to ‘HRD’ and ‘HR-proficient’. For a new sample, the Pearson correlation coefficients against the two templates were calculated, and then the correlation with the ‘HR-proficient’ template was subtracted from the correlation with the ‘HRD’ template to generate an overall HRD score.

This transcriptional signature was compared against four published signatures: a 230-gene HRD signature developed by Peng et al.(32), a 77-gene BRCA1ness signature developed by Severson et al.(31), a 70-gene signature of chromosomal instability (CIN70)(34), and a 7-gene signature of PARPi sensitivity (PARPi7)(33). Application of these signatures for comparison was also conducted using the centroid method described above. In the event of a gene in the signature not appearing in a dataset, the gene was removed from the signature and did not contribute to the correlation calculation.

#### Gene Set Enrichment Analysis

Gene set enrichment analysis was conducted using the pathfindR R package(118) and enrichR R package(119). For the pathfindR analysis, to provide relevant significance values, an ANOVA was conducted for each gene against the four BRCA-defect groups. The KEGG and Gene Ontology Biological Process gene sets were used, and default inputs were used for the remaining arguments.

### Importance Analysis using a Graph Neural Network Approach

A Graph Attention Network (GAT) was used to map gene co-expression graphs into an embedding space and to analyse its classification output in order to obtain a classification importance score for each gene, indicative of the extent to which this gene’s expression can discriminate HRD and HR-proficient samples. The pipeline consisted of following steps:

1. A weighted correlation network analysis (WGCNA)(72) method was employed to extract a gene co-expression graph involving all 228 genes in the HRD transcriptional signature across the TCGA-BRCA cohort;
2. A GAT-based feature extractor was applied to the high dimensional embeddings of the gene co-expression graphs which integrates information from neighbouring domains;
3. A simple prediction module for the classification task was implemented;
4. A gradient-based method was used to calculate gene importance scores as post-hoc explanations for model behaviour(120).

This pipeline was implemented based on the Pytorch(121) and Pytorch Geometric library(122). An Adam optimizer was used with batch size 8 and an initial learning rate of 0.002. Linear learning rate decay and early stopping were applied to avoid overfitting. A threshold of 0.4 was applied to identify ‘important’ genes for HRD and HR-proficiency classification.

### Cell-cell interaction analysis

Differential patterns of cell-cell interactivity within the tumour microenvironment of HRD and HR-proficient cells were analysed using CellphoneDB(86), which was applied to the Qian et al. and Bassez et al. cohorts(71). Tumour cells were labelled as HRD if they displayed a positive HRD score. CellphoneDB was conducted within a Conda virtual Python environment, with default parameters applied.

### Statistical analysis

Groups were compared using a two-sided Student’s t test, Wilcoxon rank-sum test or ANOVA, as appropriate. P-values were adjusted for multiple testing where appropriate using the Benjamini-Hochberg method. Graphs were generated using the ggplot2 and ggpubr R packages.

## DECLARATIONS

### Ethics, consent and permissions

All data employed in this study comply with ethical regulations, with approval and informed consent for collection and sharing already obtained by the relevant consortia where the data were obtained from (TCGA, ICGC).

### Consent for publication

Not applicable.

### Availability of data and materials

The results published here are in part based upon data generated by the TCGA Research Network (https://www.cancer.gov/tcga), ICGC (https://dcc.icgc.org/), or deposited at cBioPortal (https://www.cbioportal.org/), CCLE (https://sites.broadinstitute.org/ccle/) and GEO (https://www.ncbi.nlm.nih.gov/geo/).

All code developed for the purpose of this study can be found at the following repository: https://github.com/secrierlab/MultiscaleHRD

## Authors’ contributions

MS designed and supervised the study. DHJ developed the mutational and transcriptional classifiers and performed all analyses in bulk and single cell datasets. SP developed and applied the graph neural network classifier. JF helped co-supervise DHJ and provided valuable feedback on the analyses performed in the study.

