## Supplementary Material for "Multi-scale characterisation of homologous recombination deficiency in breast cancer"

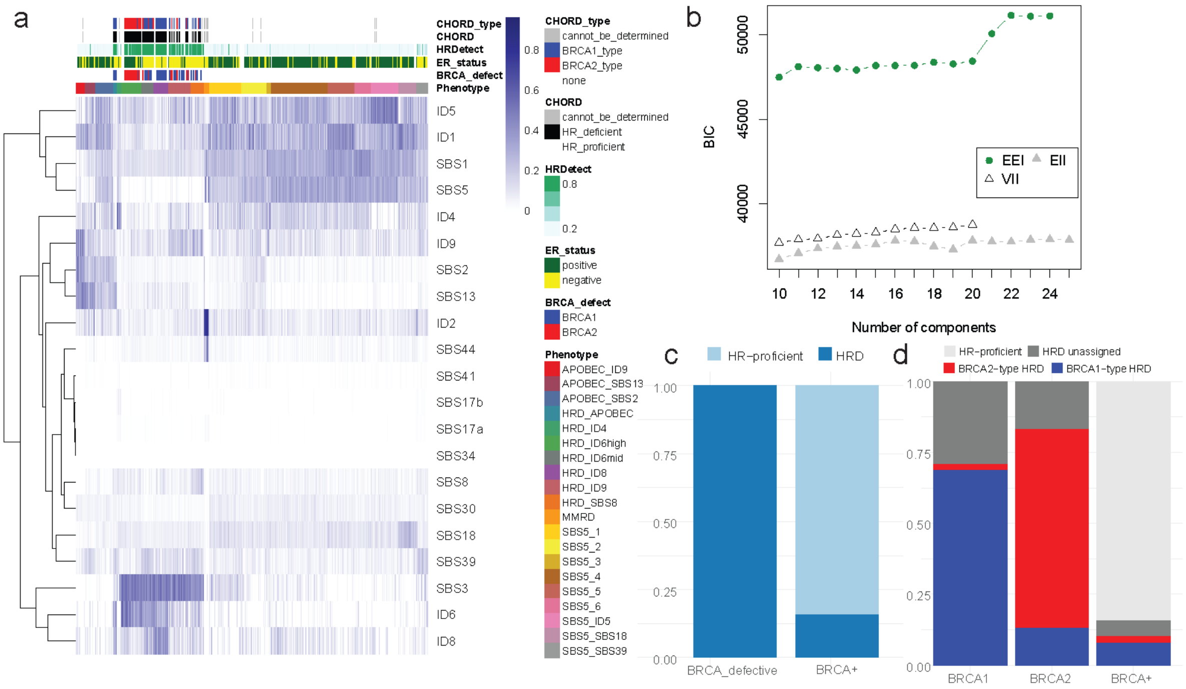


**Supplementary Figure 1. Establishment of mutational signature phenotypes in whole genome sequenced breast cancers from the ICGC cohort.** (a) Contributions of single base substitution and indel signatures. The colour represents the overall contribution of the signature to either the single base substitution or indel load of the respective sample. (b) Finite mixture modelling of the mutational signature profiles of the ICGC-BRCA cohort. The optimal clustering was determined using the Bayesian Information Criterion, with varying models tested as described by the initialisms. (c) Assigned signature phenotypes for BRCA-defective and BRCA+ samples. (d) Assigned signature phenotypes for BRCA1- and BRCA2-defective samples separately. BRCA-type HRD clusters were assigned using (a), specifying BRCA1-type HRD (HRD_APOBEC, HRD_ID6mid, HRD_ID8, HRD_SBS8), BRCA2-type HRD (HRD_ID6high), and unassigned HRD (HRD_ID4, HRD_ID9).

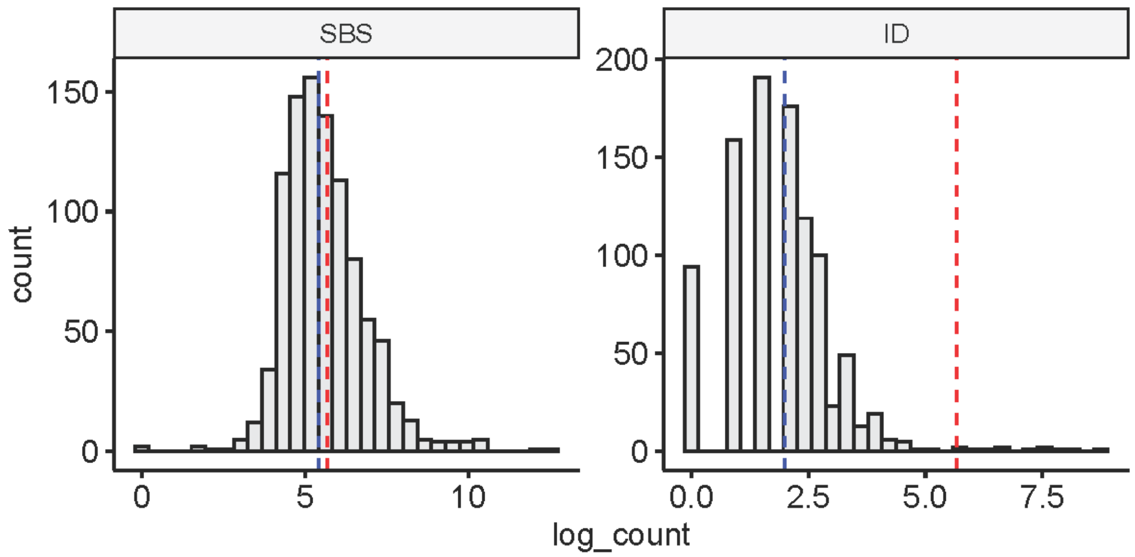


**Supplementary Figure 2. Single-base substitution (SBS) and indel (ID) loads across 968 exome sequenced breast cancers from the TCGA-BRCA cohort.** The red and blue dotted lines represent 50 mutations and the median mutational load respectively.

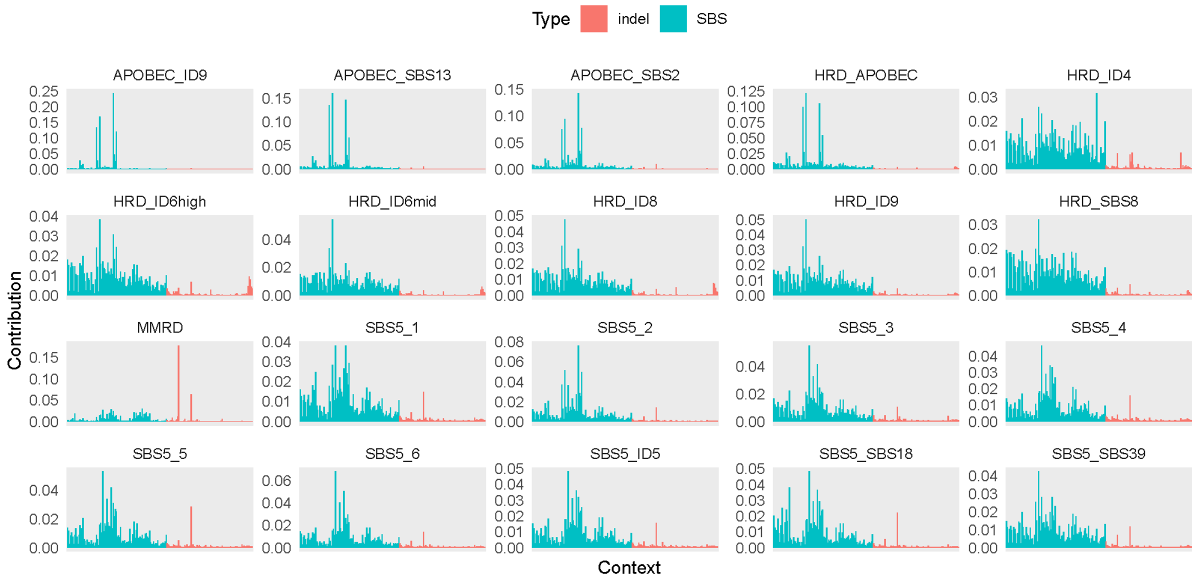


**Supplementary Figure 3. Likelihood distributions of SBS (blue) and indel (red) mutation types for each of the 20 signature phenotypes.** The likelihood distributions are calculated as the mean mutational spectrum based on associated samples from the ICGC-BRCA cohort.

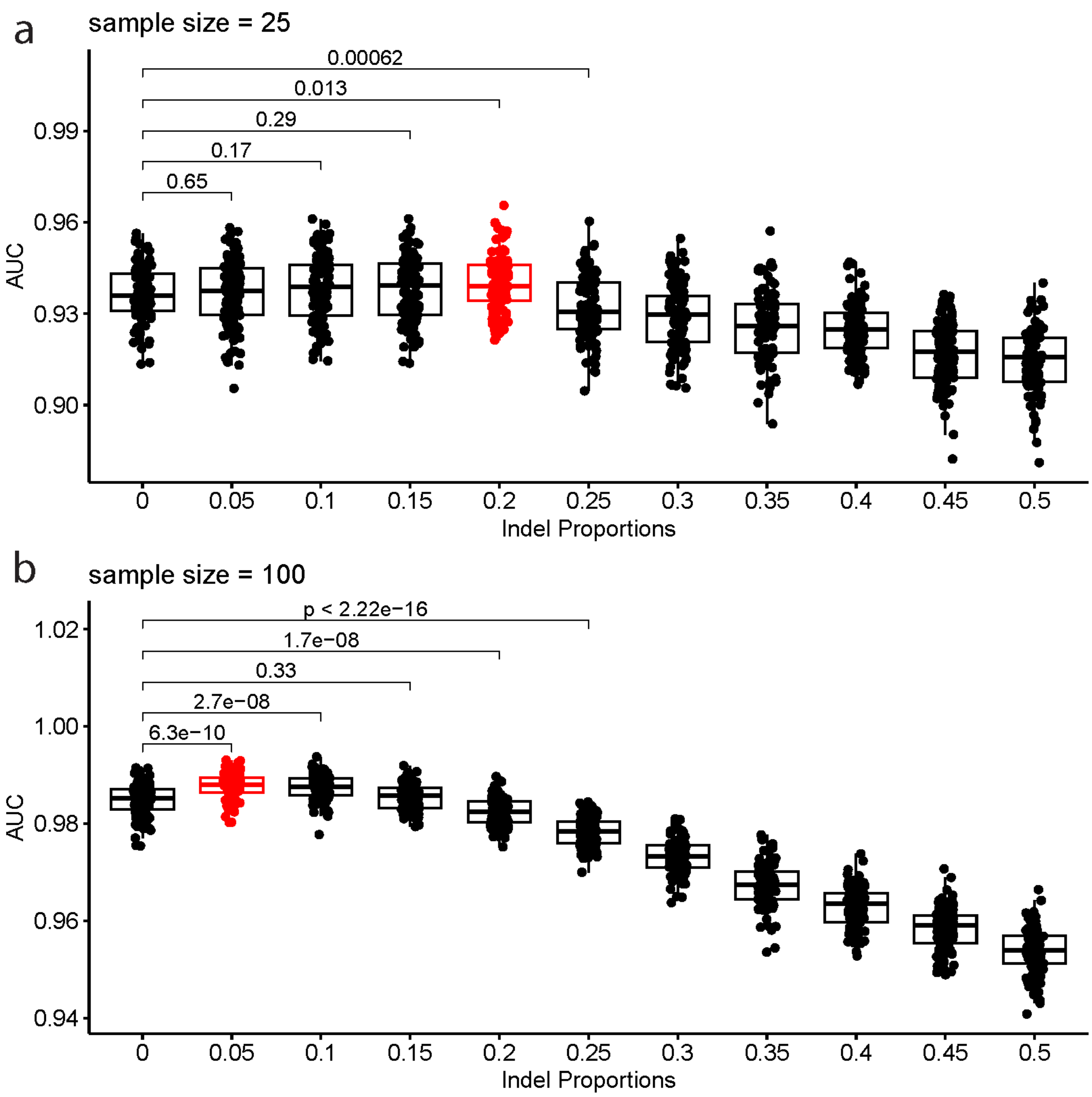


**Supplementary Figure 4. The impact of varying indel proportions on the reclassification of SBS3-enrichment in whole genome sequenced samples.** The AUC performance for SBS3 enrichment classification is displayed when considering incremental indel proportions in whole-genomes from the ICGC breast cancer cohort downsampled to (a) 25 and (b) 100 mutations. The x-axis refers to the constrained proportion of indels sampled from each sample. 100 iterations were conducted at each indel proportion between 0 and 0.5, increasing in increments of 0.05.

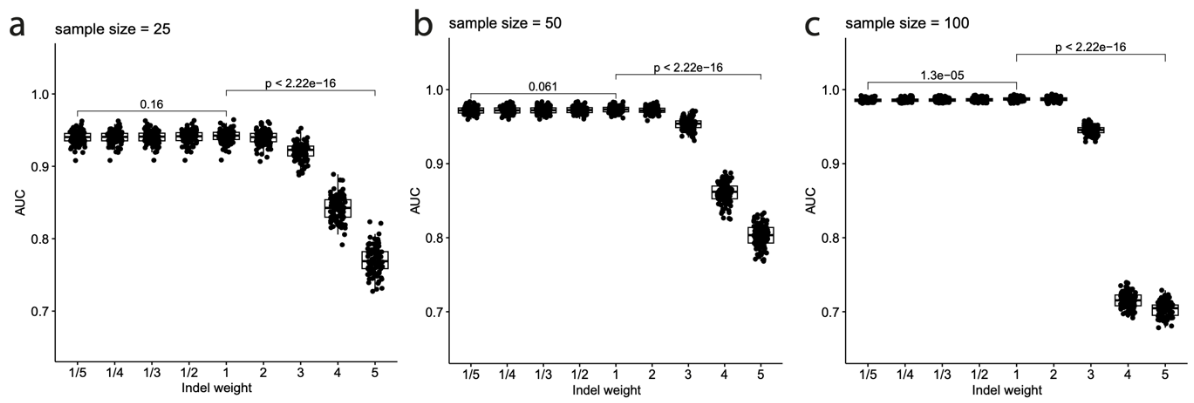


**Supplementary Figure 5. The impact of varying indel weights within likelihood distributions on reclassifying SBS3-enrichment in whole genome sequenced samples.** The AUC performance for SBS3 enrichment classification is displayed when considering different weights for indel contributions in whole-genomes from the ICGC breast cancer cohort downsampled to (a) 25 (b) 50 and (c) 100 mutations. The x-axis refers to the factor (weight) by which indel contributions are multiplied within the likelihood distributions. The AUC reflects the performance in classifying SBS3-enriched samples as such. 100 iterations were conducted at each indel weight factor.

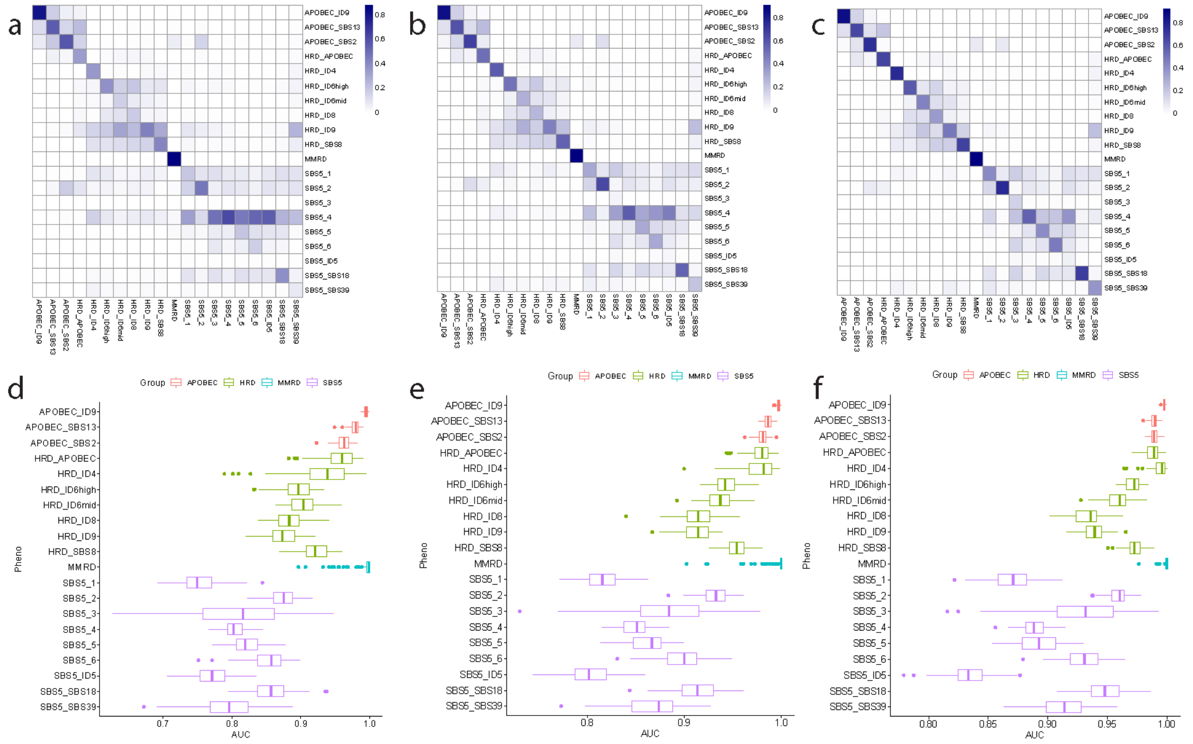

**Supplementary Figure 6. Analysis of reclassification ability for the 20 signature phenotypes at lower mutational loads.** The reclassification is performed on samples from the ICGC breast cancer cohort across 100 simulations following downsampling to (a,d) 25, (b,e) 50, and (c,f) 100 mutations. (a-c) Rows represent true signature phenotypes, columns represent assigned signature phenotypes, with the colour representing the proportion of downsampled samples across 100 simulations to be assigned to the assigned phenotype. (d-f) AUCs of reclassification for each signature phenotype across 100 simulations.

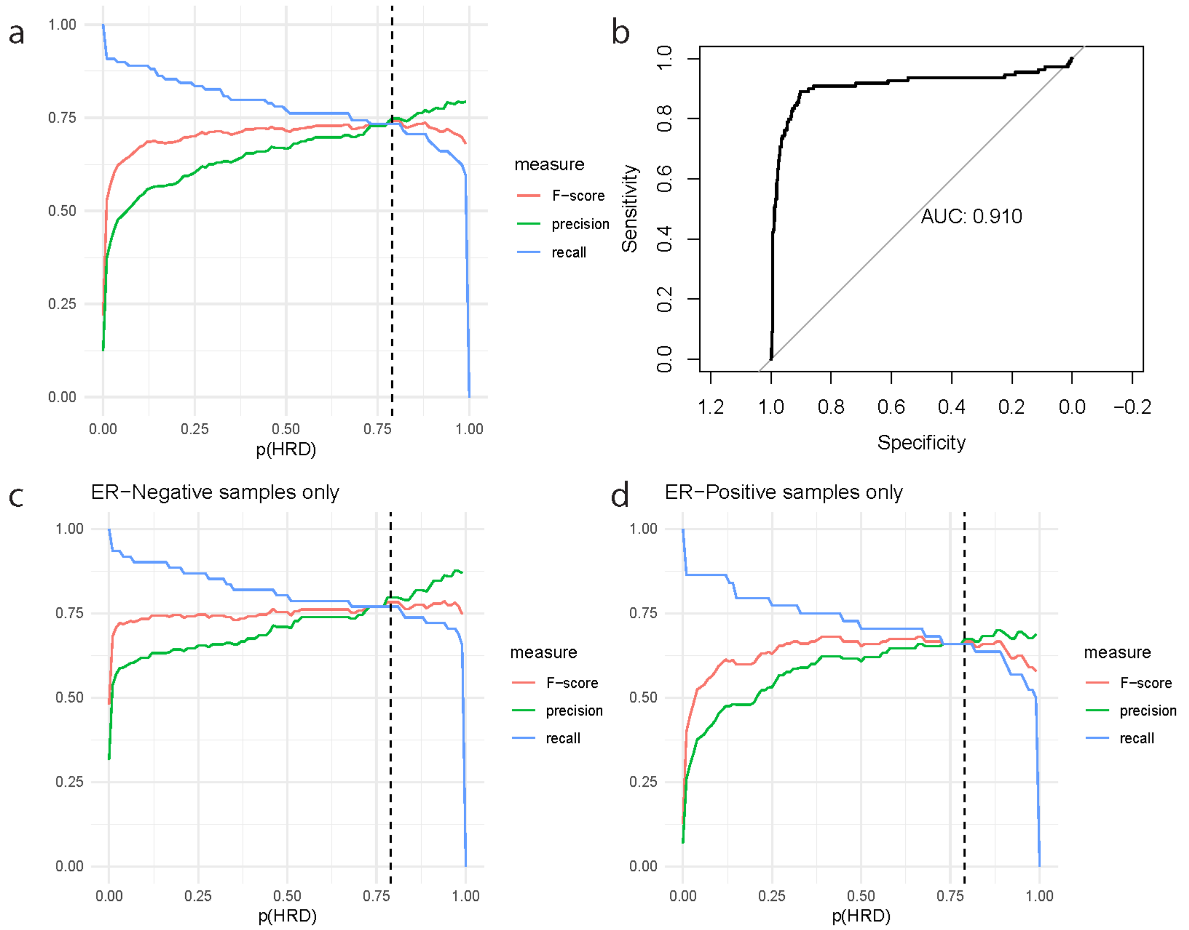


**Supplementary Figure 7. Determination of the probability threshold for HRD classification.** (a) Precision, recall, and F-scores for prediction of HR gene defects amongst the whole TCGA-BRCA cohort using the assigned probability of HRD. (b) The HRD probabilities assigned by the mutational classifier predict HR gene defects with AUC = 0.910. (c-d). Precision, recall, and F-scores for prediction of HR gene defects amongst (c) ER-negative and (d) ER-positive samples within the TCGA-BRCA cohort. The dotted black line represents p(HRD) = 0.79, the optimal threshold according to (b).

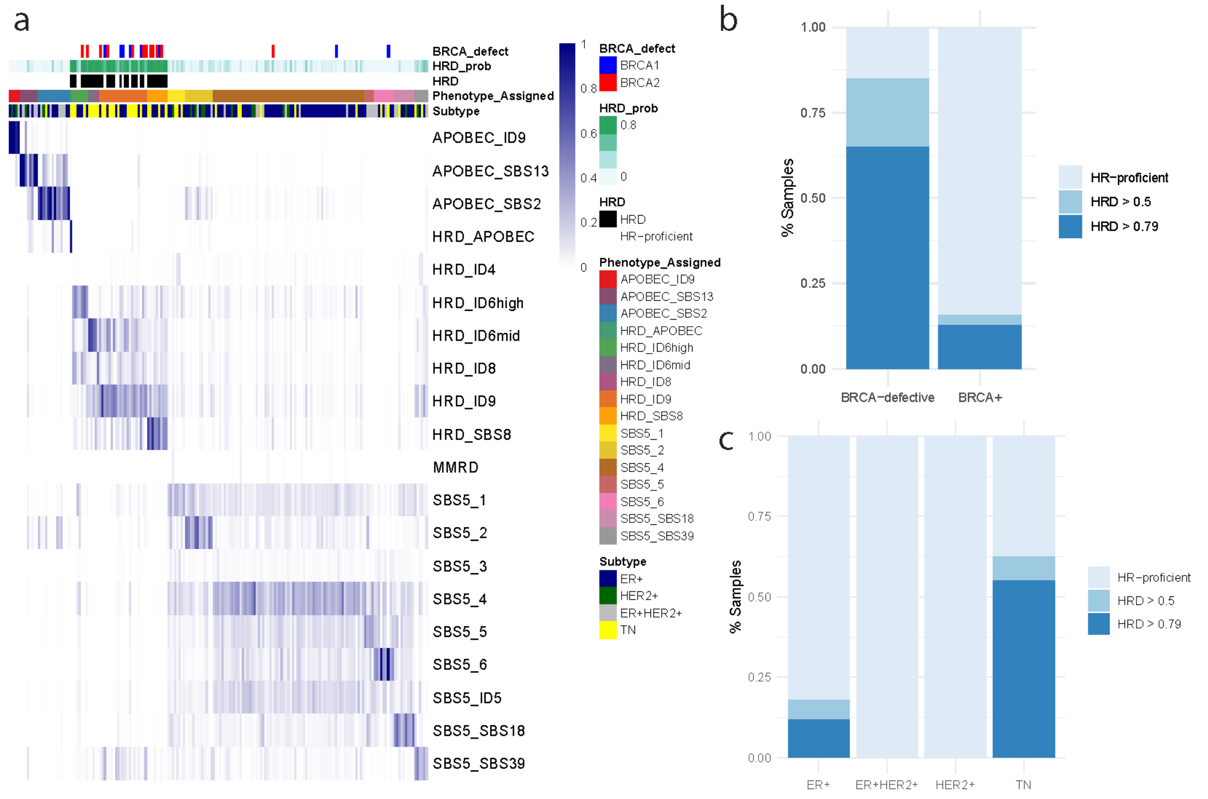


**Supplementary Figure 8. Validation of the mutation classifier in the SMC-BRCA cohort.** (a) A summary of the predictions by the mutation classifier, with rows representing samples and the colour representing the probability of assignment to the respective signature phenotype. 'Phenotype_Assigned' refers to the phenotype with the highest probability of assignment, and 'HRD_prob' is the sum of the probabilities of assignment across the seven HRD-associated phenotypes. (b) The 20 BRCA-defective samples are substantially more likely to be assigned as HRD compared with BRCA+ samples. 13/20 display an HRD probability greater than 0.79, and an additional four samples display an HRD probability between 0.5 and 0.79. (c) HRD assignment across breast cancer subtypes, with triple-negative (TN) samples enriched for HRD.

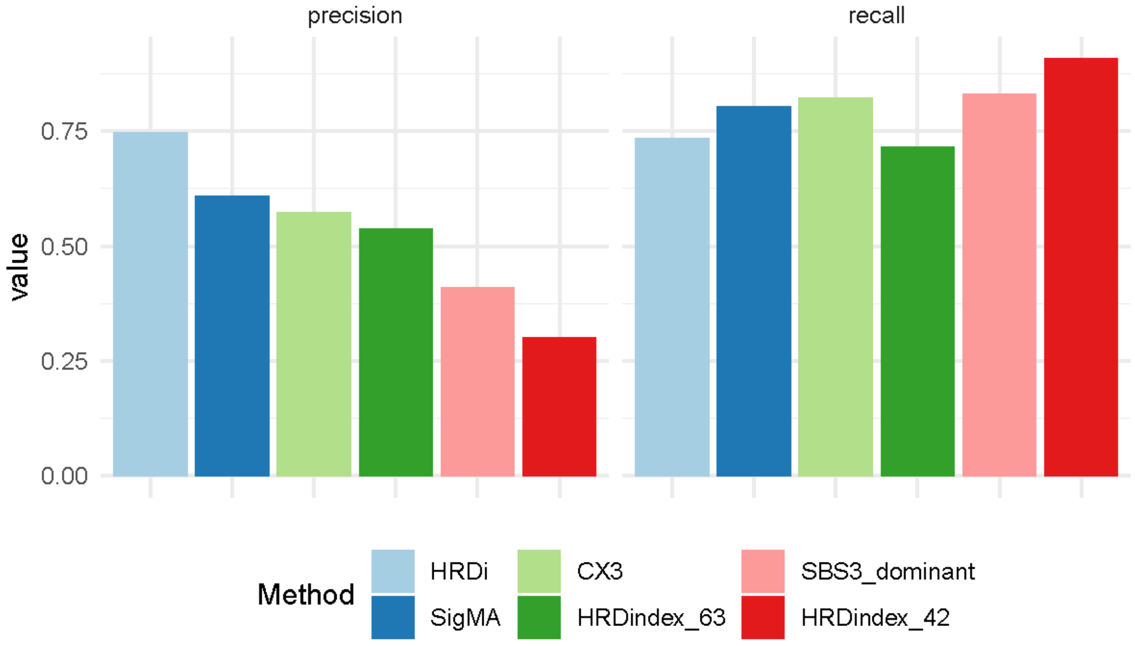


**Supplementary Figure 9. Precision and recall across HRD classifiers.** ‘HRDi’ refers to our likelihood-based classifier with a probability threshold of 0.79. ‘SigMA’ refers to SigMA application using their stringent threshold. ‘CX3’ refers to the CX3 copy number signature, in which samples with CX3 as their most prevalent signature are assigned as HRD. Similarly, ‘SBS3_dominant’ refers to samples for which SBS3 is the dominant single base substitution signature. ‘HRDindex_63’ and ‘HRDindex_42’ refer to the Myriad HRD score, applying thresholds for HRD classification of 63 and 42 respectively.

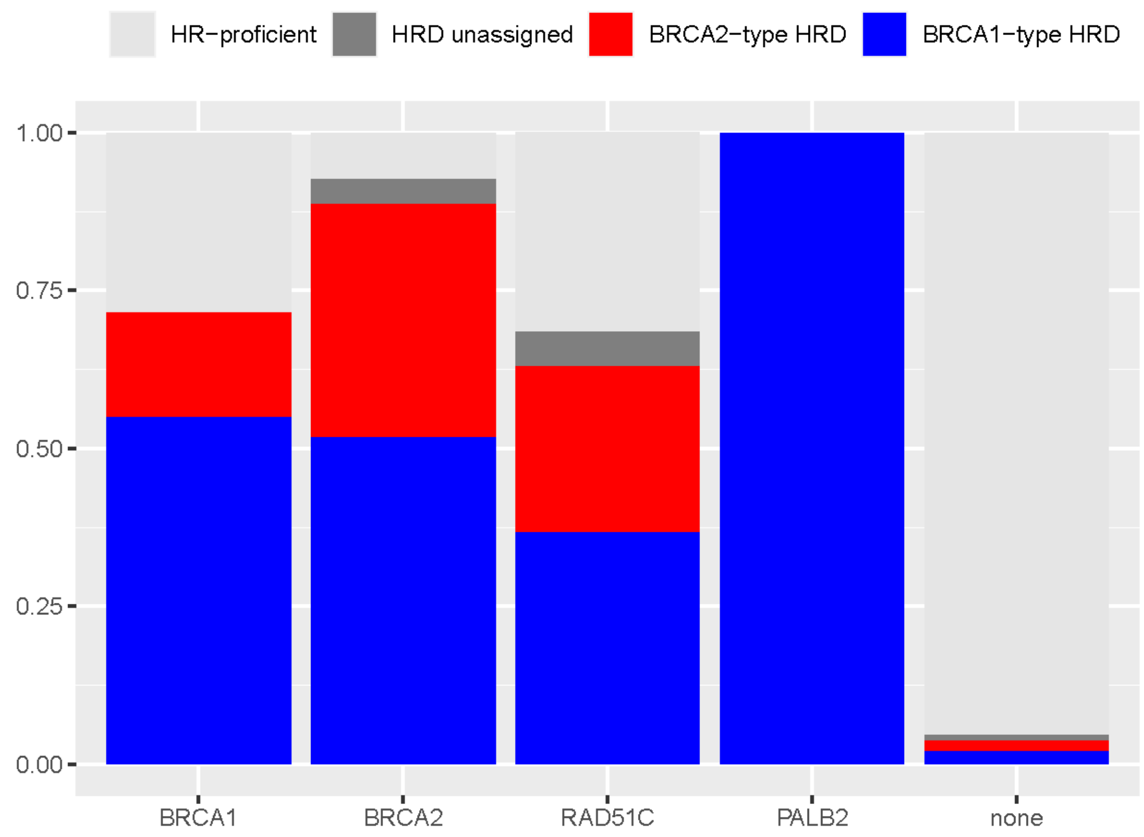


**Supplementary Figure 10. BRCA type-specific classification in TCGA.** Samples are labelled as BRCA1-type HRD (HRD_APOBEC, HRD_ID6mid, HRD_ID8, HRD_SBS8), BRCA2-type HRD (HRD_ID6high), HRD_unassigned (HRD_ID4, HRD_ID9), or HR-proficient, and are grouped according to the HR gene defects which they harbour: BRCA1 (n=60), BRCA2 (n=27), RAD51C (n=19), PALB2 (n=3), none (n=772).

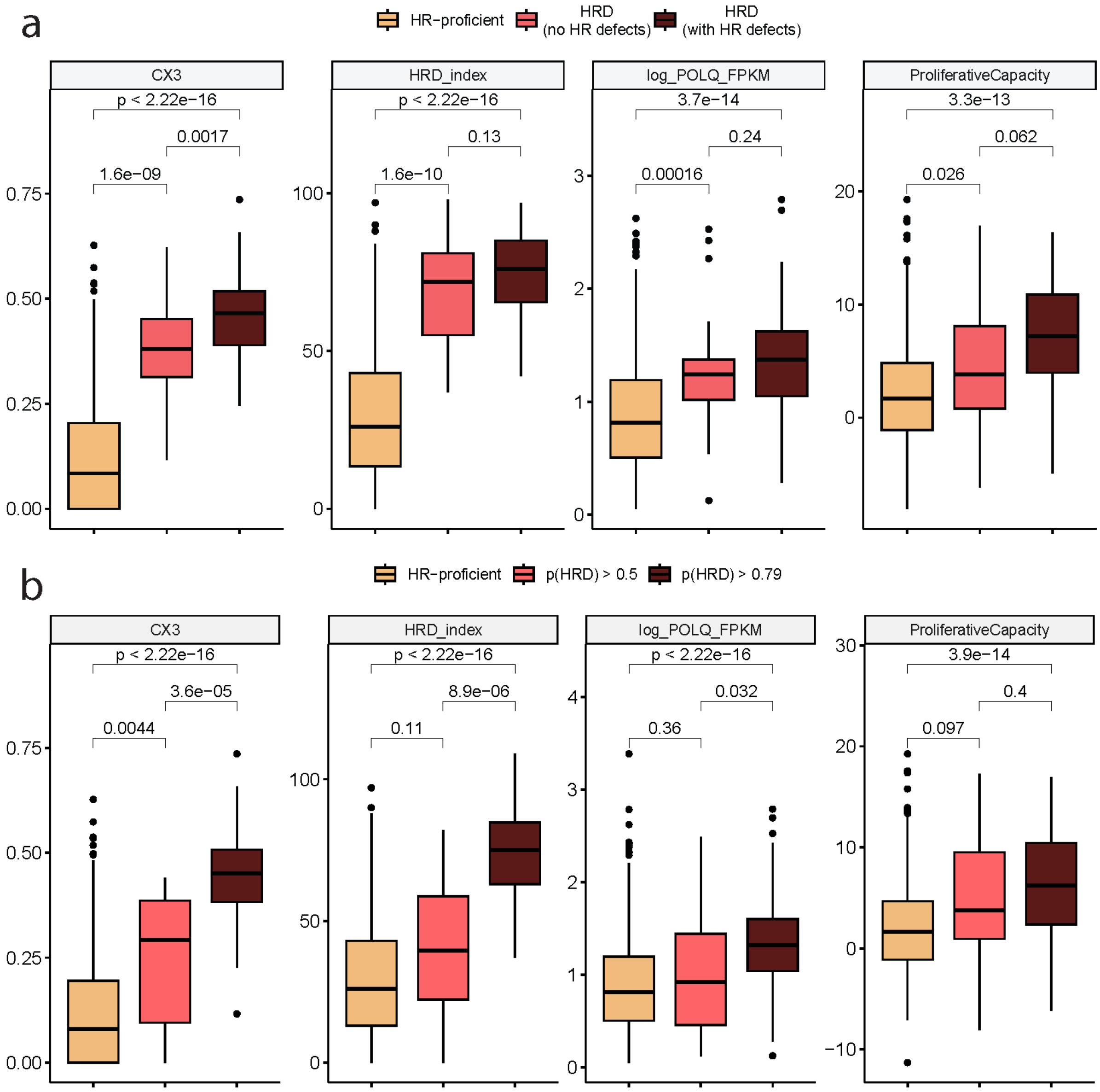


**Supplementary Figure 11. Key hallmarks of HRD across varying HRD groups and categories.** (a) Amongst HRD samples, key hallmarks broadly do not differ depending on defects in HR genes, especially in comparison with HR-proficient samples. (b) Samples with an HRD probability between 0.5 and 0.79 are more similar in terms of key HRD hallmarks to HR-proficient samples than samples with an HRD probability greater than the established threshold of 0.79.

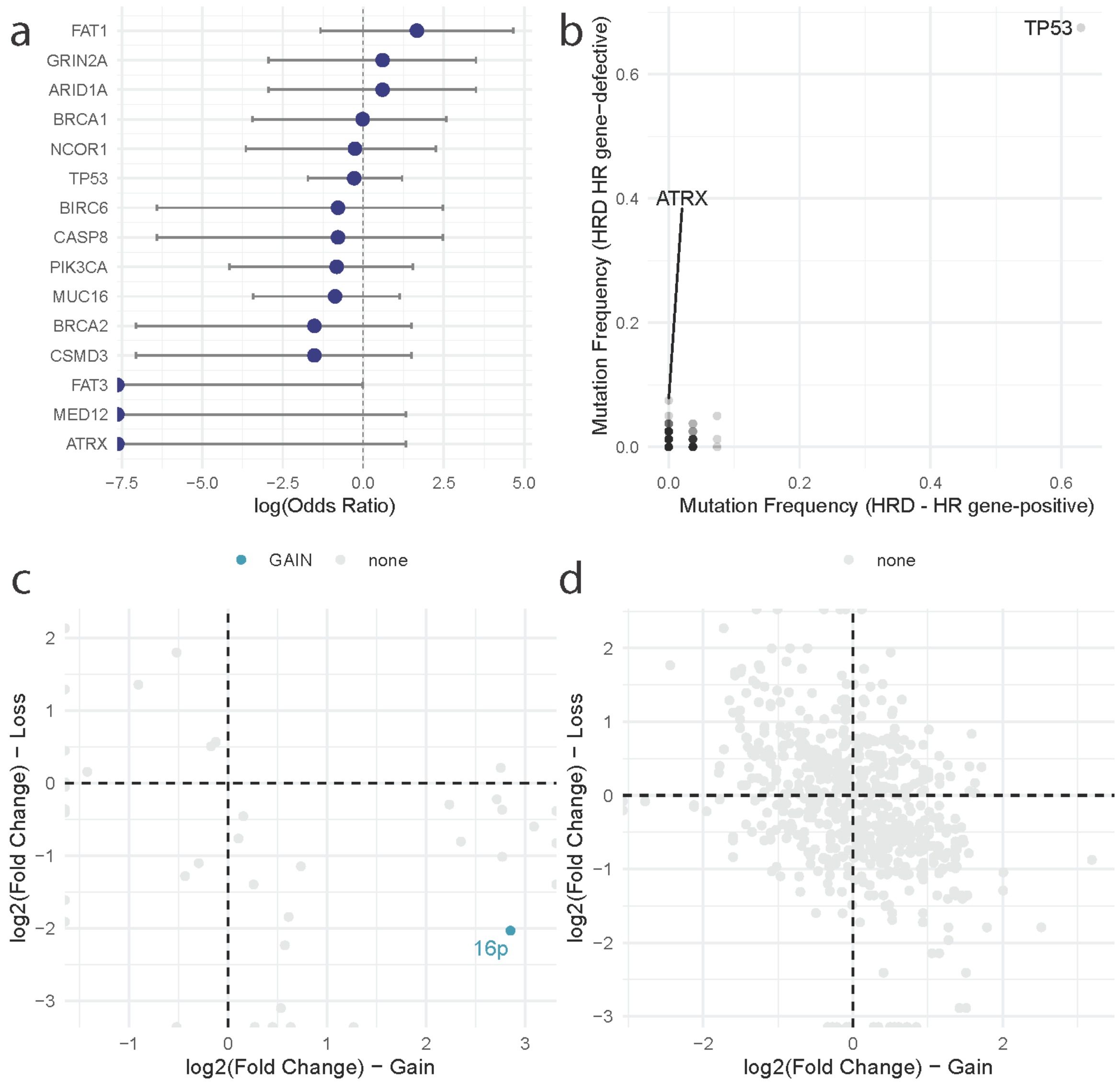


**Supplementary Figure 12. Cancer driver gene alterations between HRD HR gene-proficient and HR gene-defective samples.** (a) Odds ratios reflecting mutation enrichment (>0) or depletion (<0) of cancer driver genes mutated in >5% samples (according to a Fisher's exact test). (b) Comparison of frequency of nonsynonymous cancer driver gene mutations across HRD samples depending on their HR gene-defect status. Every dot corresponds to a gene. Only notable genes are labelled. (c) Enrichment of chromosome arm alterations in HRD samples comparing HR gene-positive against HR gene-defective samples. Positive values indicate an enrichment of gains/losses in HR gene-positive samples, whilst negative values indicate enrichment in HR gene-defective samples. Arms are coloured for whether they are enriched for gains (blue) or no significant enrichment (grey). A single significant enrichment is highlighted (16p loss). (d) Enrichment of cancer driver gene gains and losses, following the same labelling as (c). No significant enrichments are observed.

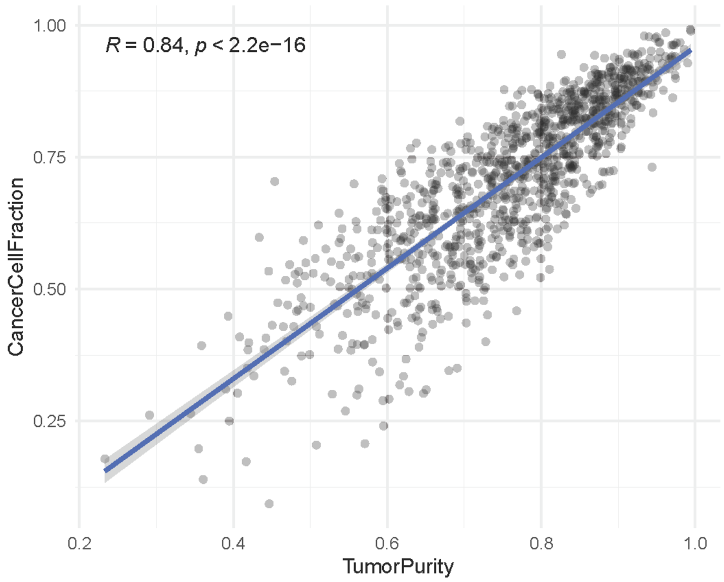


**Supplementary Figure 13. Concordance between TME-adjusted cancer cell fractions and tumour purity.** The estimated cancer cell fraction of the TCGA-BRCA cohort according to BayesPrism is compared against estimates of tumour purity inferred from the genomic data available via the TCGAbiolinks R package.

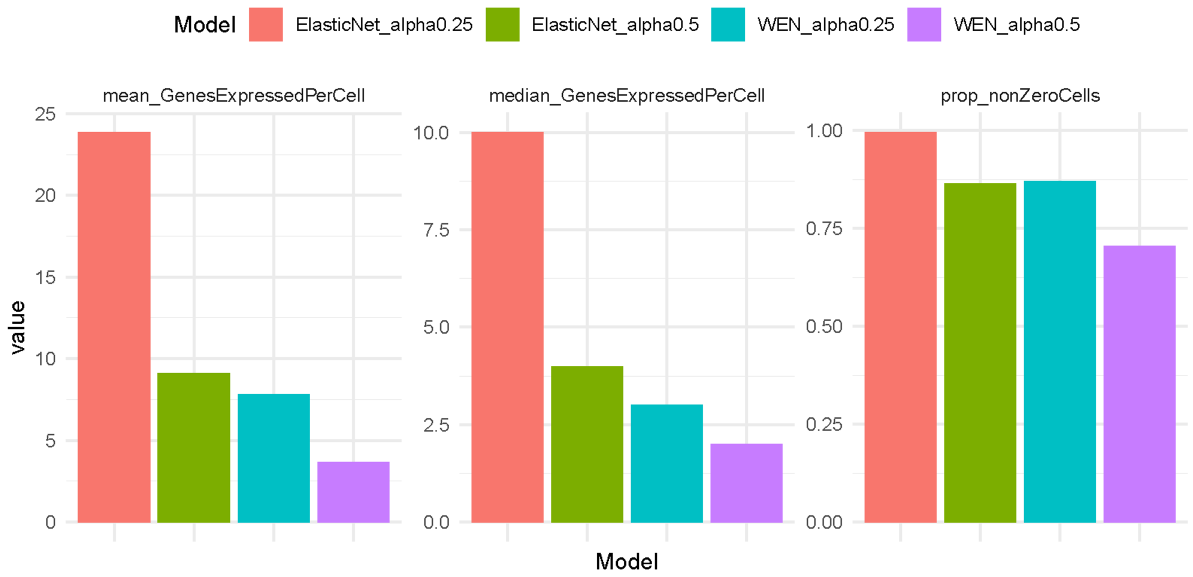


**Supplementary Figure 14. Comparisons of single cell expression for multiple signatures generated by regularised logistic regression**. Expression statistics are calculated using the Qian et al. single-cell RNA-seq breast cancer cohort. Four signatures were initially developed, without and with weighted elastic net ('ElasticNet' and 'WEN' respectively) and with regularisation penalties of 0.25 ('alpha0.25') and 0.5 ('alpha0.5'). The measures refer to the mean number of genes expressed in the respective signature per cell (left), the median number of genes expressed in the respective signature per cell (center), and the proportion of cells within the cohort which express at least one gene within the respective signature (right). The Elastic Net with 0.25 regularisation penalty outperforms other methods.

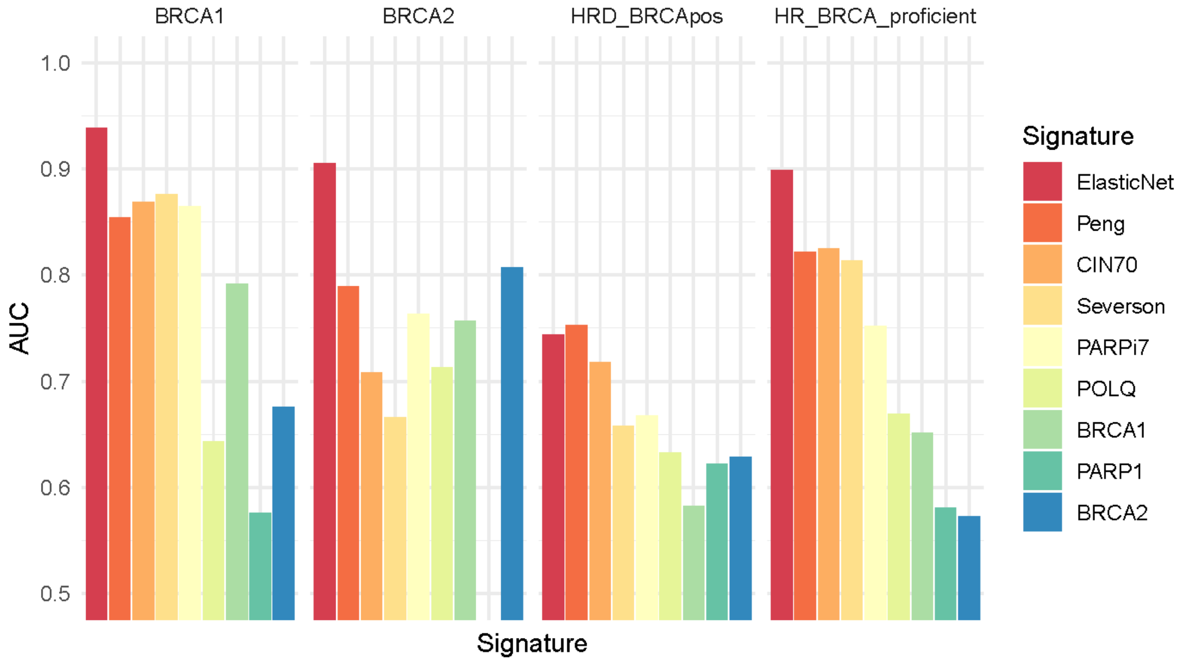


**Supplementary Figure 15. Accuracy of transcriptional signatures for predicting BRCA1-defects, BRCA2-defects, BRCA-positive HRD, and HR/BRCA-proficiency within the TCGA-BRCA testing cohort.** 'ElasticNet' refers to the in-house 228-gene signature developed here. 'Peng', 'CIN70', 'Severson', and 'PARPi7' refer to alternative HRD signatures. 'POLQ', 'BRCA1', 'PARP1', and 'BRCA2' refer to FPKM-normalised expression of the respective genes.

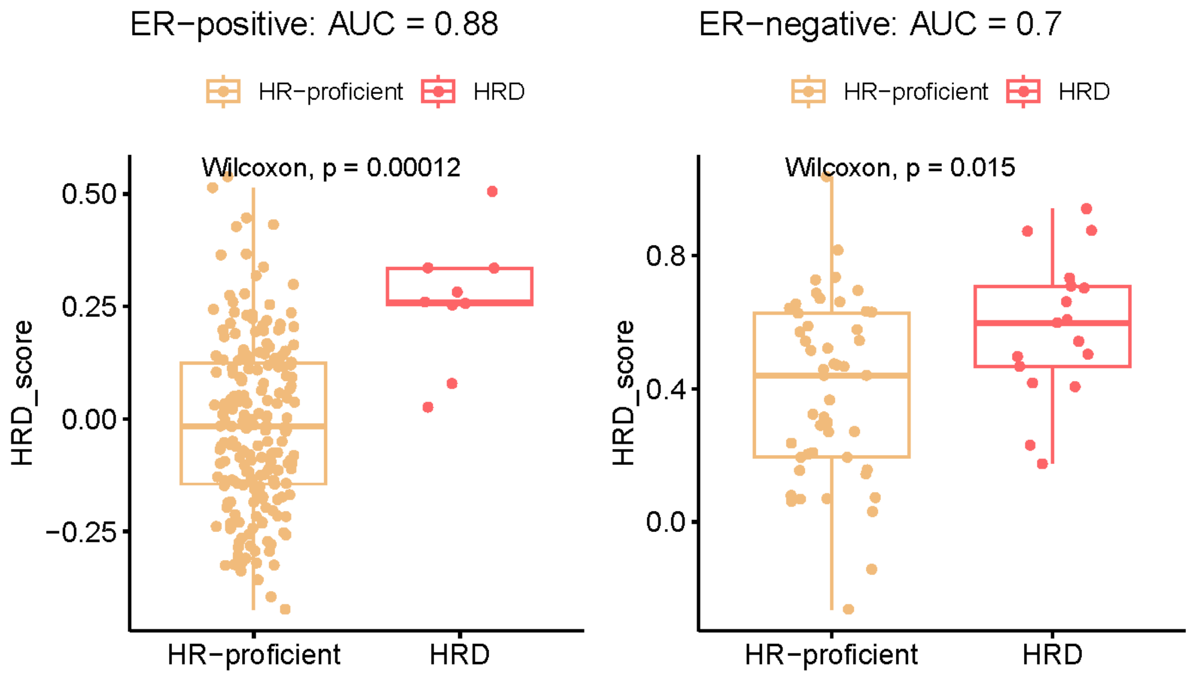


**Supplementary Figure 16.** Comparison of transcriptional HRD scores between HRD and HR-proficient samples within the TCGA-BRCA testing cohort, following separation of samples by ER status.

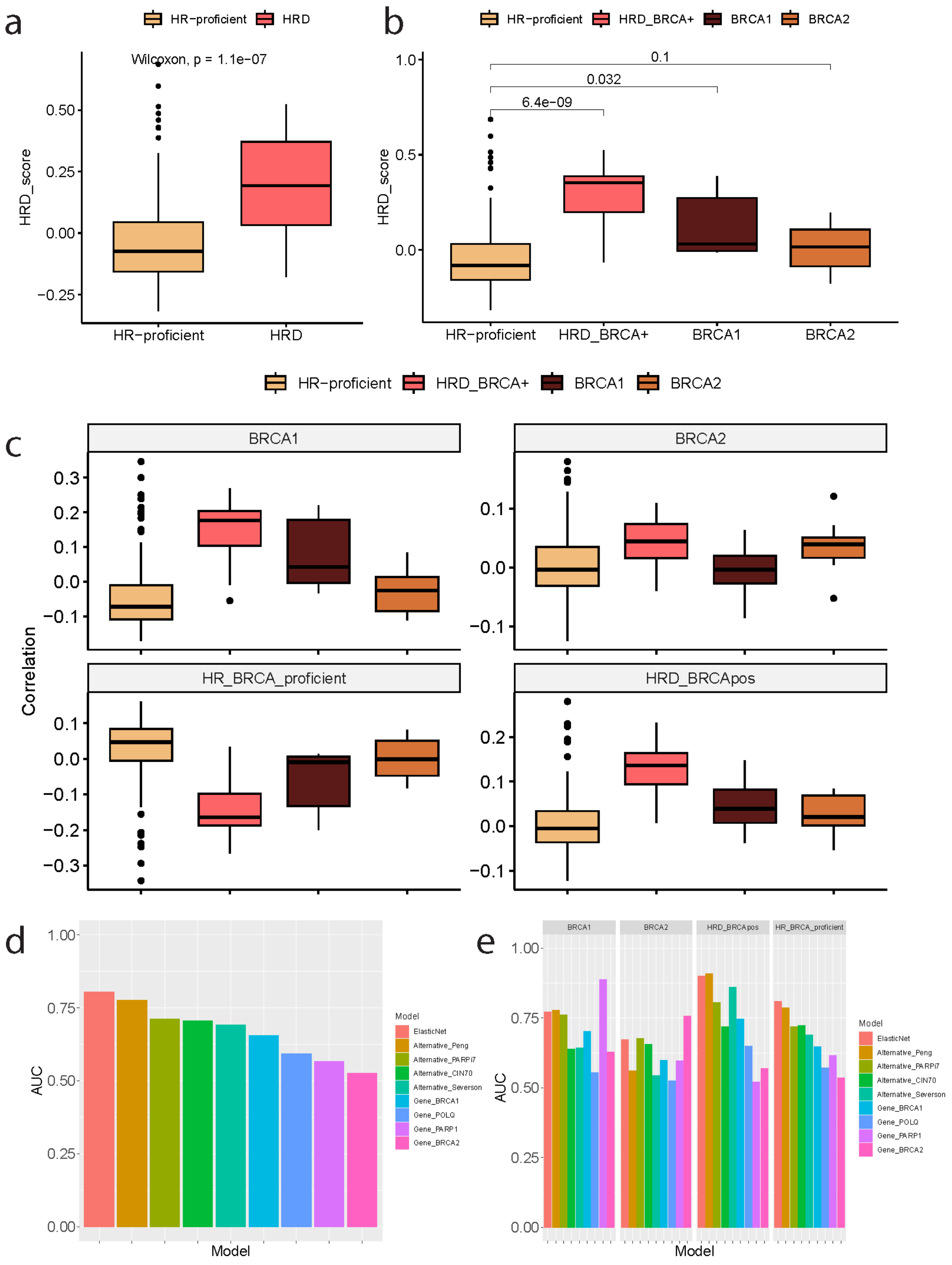


**Supplementary Figure 17. Application of the 228-gene HRD transcriptional signature to the SMC-BRCA validation cohort.** Within the cohort, this transcriptional signature is significantly increased within (a) HRD samples, and (b) BRCA1-defective and BRCA-positive HRD samples. (c) BRCA-specific transcriptional scores. (d) Accuracy of HRD classification across transcriptional signatures and HRD markers. (e) Accuracy of BRCA-specific HRD classification across transcriptional signatures and HRD markers. 'ElasticNet' refers to the 228-gene signature introduced here.

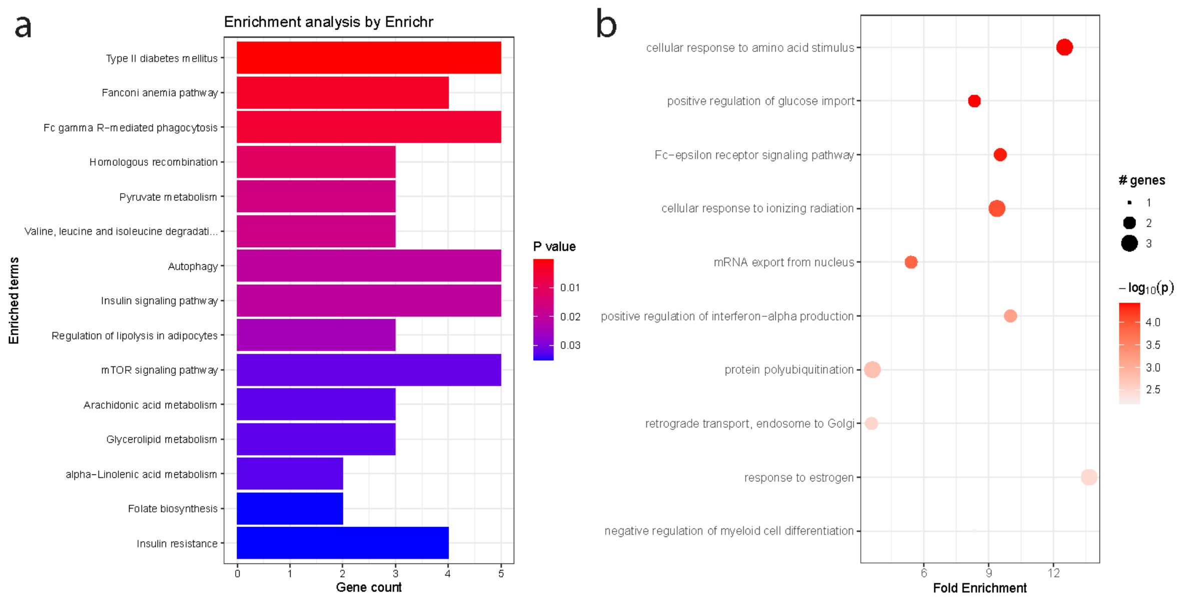


**Supplementary Figure 18. Gene Set Enrichment Analysis for the 228-gene HRD transcriptional signature**. Enrichment results are shown according to (a) enrichR using the KEGG database, and (b) pathfindR using the Gene Ontology Biological Processes resource.

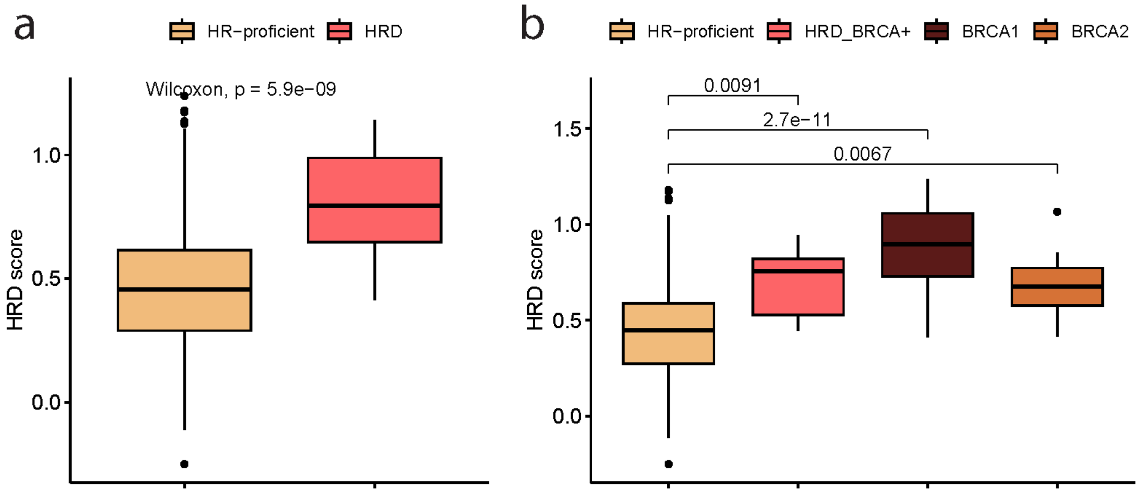


**Supplementary Figure 19. Performance of the reduced 26-gene HRD transcriptional signature in the TCGA-BRCA testing cohort.** The HRD score is compared between (a) HRD and HR-proficient samples, and (b) BRCA-specific HRD categories.

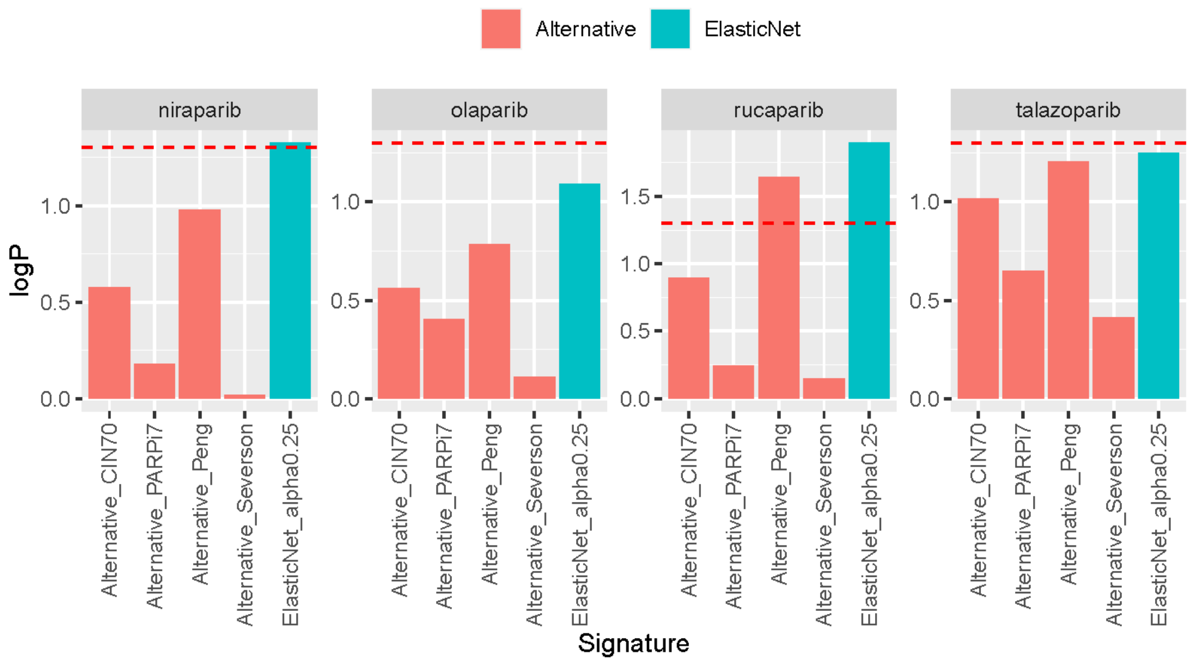


**Supplementary Figure 20. Comparison of HRD transcriptional signatures in predicting PARP inhibitor sensitivity in CCLE**. Signature performance is measured by the significance of correlation, on a negative-log10 scale, between the respective signature score and PRISM measures of PARP inhibitor sensitivity. The dotted red line represents a significance level p = 0.05, with bars exceeding this threshold demonstrating a greater significance of correlation. ‘ElasticNet’ represents the 228-gene HRD signature introduced in this study.

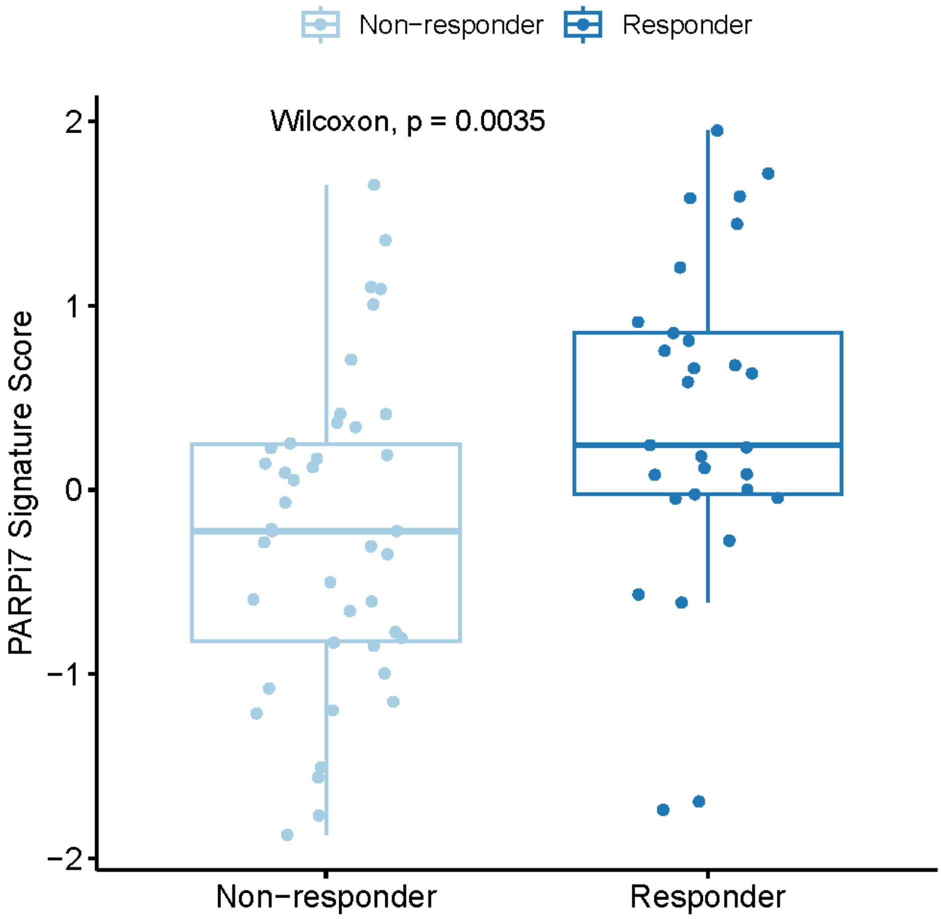


**Supplementary Figure 21. Performance of the PARPi7 transcriptional signature for predicting response to olaparib/durvalumab in patients within the treatment arm of the I-SPY2 trial.** The PARPi7 signature score is compared between responders and non-responders.

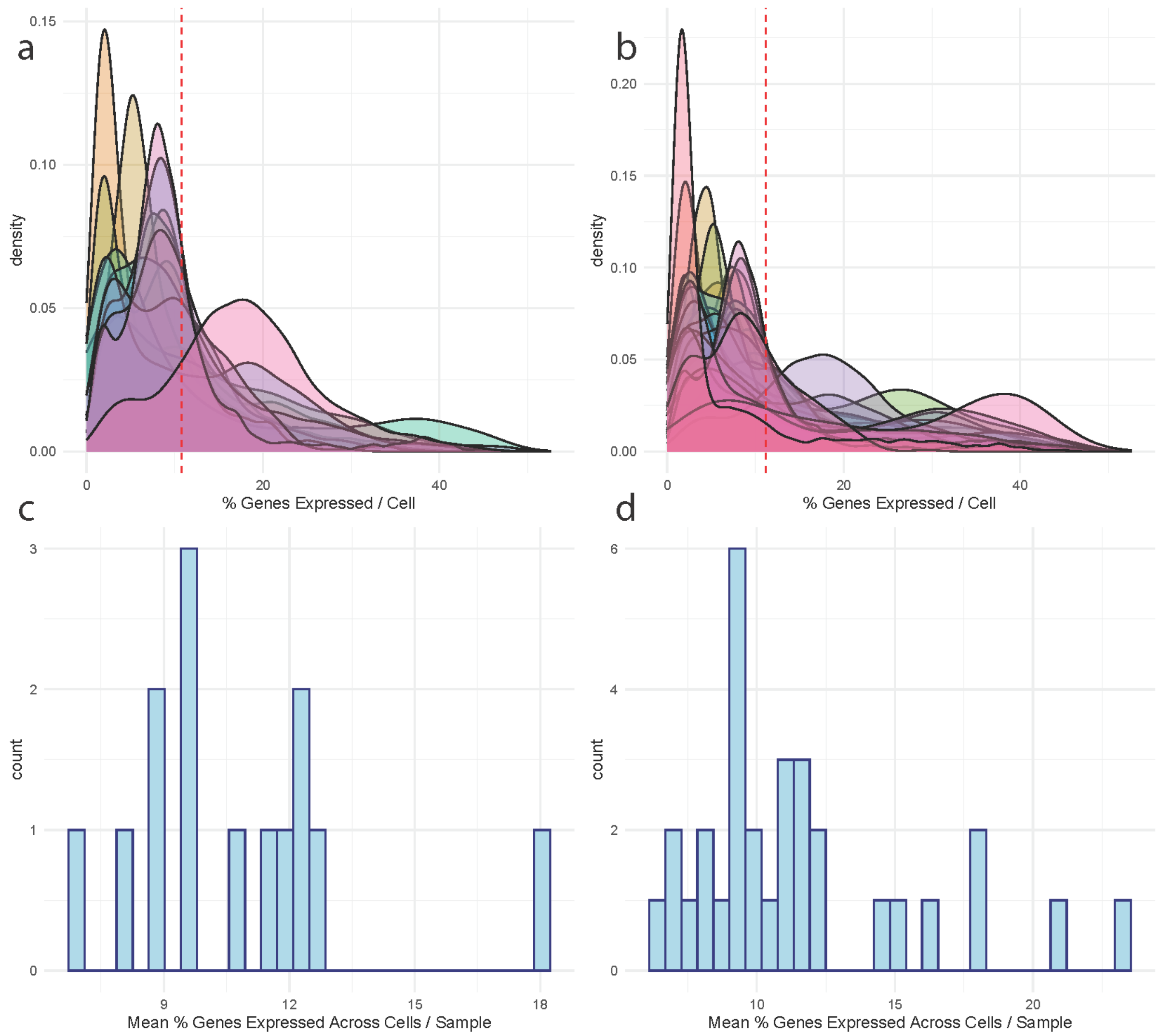


**Supplementary Figure 22. Signature expression across single-cell RNA-seq breast cancer cohorts.** (a-b) Density plot displaying the percentage of the 228 genes in the HRD transcriptional signature expressed per cell, separated by samples, across the (a) Qian et al. and (b) Bassez et al. cohorts. The dotted red lines represent the mean percentage of genes expressed per cell across the entire cohort. (c-d) Summaries of the mean percentage of the 228 genes expressed per cell across each sample within the (c) Qian et al. and (d) Bassez et al. cohorts.

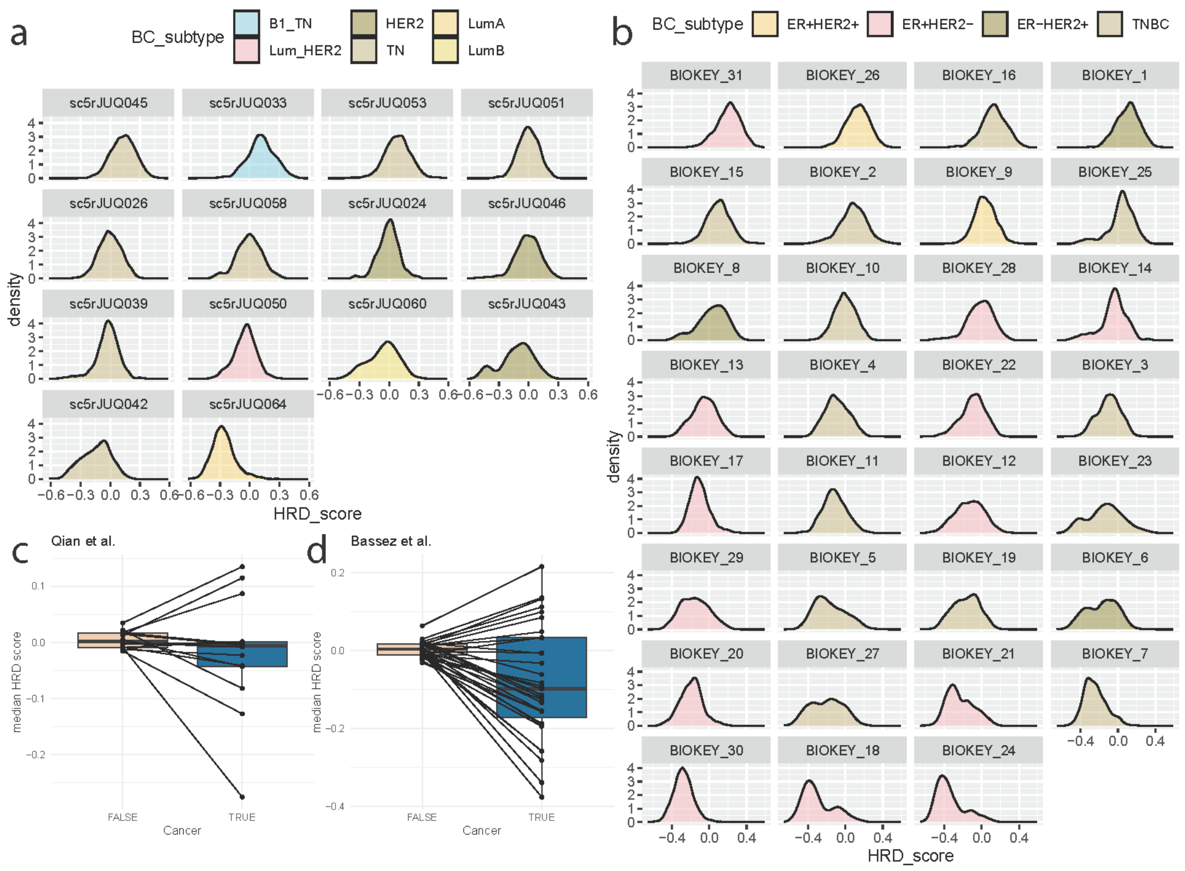


**Supplementary Figure 23. Transcriptional signals of HRD across cancer cells and the tumour microenvironment in single-cell RNA-seq breast cancer cohorts.**

(a,b) Distributions of HRD scores across individual samples from the (a) Qian et al. and (b) Bassez et al. breast cancer cohorts, coloured by assigned breast cancer subtype. (c,d) Differences in median HRD scores from cells within non-cancer and cancer cells, across samples from the (c) Qian et al. and (d) Bassez et al. cohorts. Breast cancer subtypes for Qian et al. are as follows: ‘B1 TN’ = BRCA1-defective Triple negative, ‘Lum HER2’ = Luminal-HER2+, ‘HER2’ = HER positive, ‘TN’ = Triple negative, ‘LumA’ = Luminal A-like, ‘LumB’ = Luminal B-like.

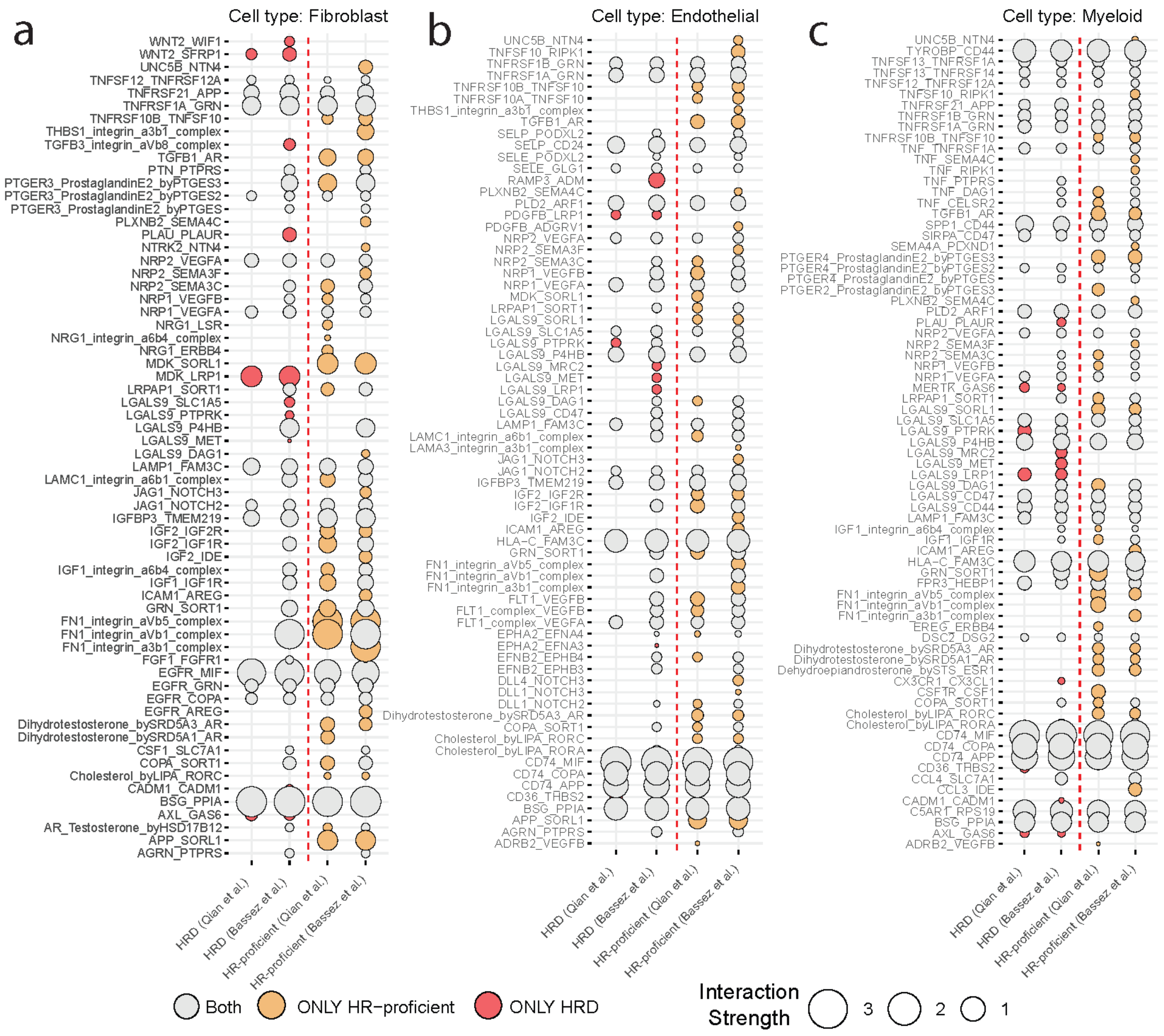


**Supplementary Figure 24. Significant ligand-receptor interactions between various immune/stromal cells and cancer cells.** Interactions between cells int the TME (as sources) and cancer cells (as targets) are displayed based on the HRD/HR-proficent status of the tumour cells in the Qian et al. and Bassez et al. single-cell RNA-seq cohorts according to CellphoneDB. Respective source cell types are (a) fibroblasts, (b) endothelial cells, and (c) myeloid cells. The strength of the interaction is defined by the size of the circle. Unique interactions for HRD cells are coloured in red, and those for HR-proficient cells are coloured in yellow.
